# The RNA binding protein Nab2 regulates splicing of the RhoGEF *trio* transcript to govern axon and dendrite morphology

**DOI:** 10.1101/2024.04.05.588324

**Authors:** Carly L. Lancaster, Pranav S. Yalamanchili, Jordan N. Goldy, Sara W. Leung, Anita H. Corbett, Kenneth H. Moberg

## Abstract

The *Drosophila* RNA binding protein (RBP) Nab2 acts in neurons to regulate neurodevelopment and is orthologous to the human intellectual disability-linked RBP, ZC3H14. Nab2 governs axon projection in mushroom body neurons and limits dendritic arborization of class IV sensory neurons in part by regulating splicing events in ~150 mRNAs. Analysis of the *Sex-lethal* (*Sxl*) mRNA revealed that Nab2 promotes an exon-skipping event and regulates m^6^A methylation on *Sxl* pre-mRNA by the Mettl3 methyltransferase. Mettl3 heterozygosity broadly rescues *Nab2^null^* phenotypes implying that Nab2 acts through similar mechanisms on other RNAs, including unidentified targets involved in neurodevelopment. Here, we show that Nab2 and Mettl3 regulate the removal of a 5’UTR intron in the *trio* pre-mRNA. Trio utilizes two GEF domains to balance Rac and RhoGTPase activity. Intriguingly, an isoform of Trio containing only the RhoGEF domain, GEF2, is depleted in *Nab2^null^* nervous tissue. Expression of Trio-GEF2 rescues projection defects in *Nab2^null^*axons and dendrites, while the GEF1 Rac1-regulatory domain exacerbates these defects, suggesting Nab2-mediated regulation Trio-GEF activities. Collectively, these data identify Nab2-regulated splicing as a key step in balancing Trio GEF1 and GEF2 activity and show that Nab2, Mettl3, and Trio function in a common pathway that shapes axon and dendrite morphology.

**Significance Statement:**

- *Drosophila* Nab2, ortholog of the human RBP ZC3H14 mutated in inherited intellectual disability, acts through unknown RNA targets to control axon and dendrite morphology.
- This study shows that Nab2 and the Mettl3 methyltransferase guide splicing of *trio* mRNA, which encodes a conserved GEF-domain protein. Intron retention in *trio* mRNA leads to an imbalance in levels of two Trio GEF domains in Nab2-deficient neurons and restoring this balance rescues neuronal defects.
- The authors conclude that Nab2 control of *trio* splicing is required to pattern axon and dendrite growth and suggests that ZC3H14 may play a similar role in the vertebrate brain.

## INTRODUCTION

RNA binding proteins (RBPs) associate with nascent RNA transcripts and govern expression via a multitude of mechanisms, including regulation of splicing, polyadenylation, nuclear export, translation, and stability (Maniatis and Reed, 2002; McKee and Silver, 2007; Corbett, 2018; Corley *et al*., 2020). These RNA-RBP interactions are particularly important in highly specialized cells such as neurons which require fine-tuned spatiotemporal control of gene expression to ensure proper development and function of the nervous system (Bardoni *et al*., 2012; Santoro *et al*., 2012; Conlon and Manley, 2017; Thelen and Kye, 2019; Gebauer *et al*., 2021). The importance of RBP function in neurons is highlighted by the prevalence of neurodevelopmental diseases that have been linked to defects in RBPs, leading to aberrant processing of RNAs encoding neurodevelopment factors (Cooper *et al*., 2009; Pak *et al*., 2011; Bardoni *et al*., 2012; Darnell and Richter, 2012; Gross *et al*., 2012; Santoro *et al*., 2012; Edens *et al*., 2015; Agrawal *et al*., 2019). Intriguingly, many of these RBPs are ubiquitously expressed and have roles in relatively common RNA processing mechanisms (Barbe *et al*., 1996; Franke *et al*., 1996; Brais, 2003; Lage *et al*., 2008; Kolb and Kissel, 2011; Pirozzi *et al*., 2011). Therefore, defining roles for these RBPs in neurons has become key to understanding why they are linked to neurological disease.

One important family of post-transcriptional regulatory proteins consists of polyadenosine binding proteins (Pabs) (Kelly *et al*., 2010). Conventional Pab family members bind polyadenosine RNA via RNA recognition motifs (RRMs) and modulate a multitude of RNA processing events such as splicing, export, polyadenylation, translation, and stability (Banerjee *et al*., 2013; Goss and Kleiman, 2013; Wigington *et al*., 2014). Another less well-studied group of Pabs utilize zinc finger (ZnF) domains to bind specific RNA motifs and modulate downstream processing events (Kelly *et al*., 2007; Kelly *et al*., 2010; Kelly *et al*., 2014). One such ZnF Pab termed <u>z</u>inc finger <u>C</u>ys-<u>C</u>ys-<u>C</u>ys-<u>H</u>is-type containing <u>14</u> (ZC3H14; also termed MSUT2) is expressed ubiquitously and binds tracts of polyadenosine RNA with high affinity via tandem ZnF domains (Kelly *et al*., 2007; Leung *et al*., 2009; Kelly *et al*., 2010; Kelly *et al*., 2014; Wheeler *et al*., 2019). Despite ubiquitous expression, mutations in human *ZC3H14* cause a form of inherited non-syndromic autosomal recessive intellectual disability, which implies a specific requirement for ZC3H14 in the developing brain (Pak *et al*., 2011; Kelly *et al*., 2012).

The ZC3H14 protein is evolutionarily conserved among eukaryotes and has been studied in *Mus musculus* (*Zc3h14*) (Guthrie *et al*., 2011; Pak *et al*., 2011; Soucek *et al*., 2016), *Caenorhabditis elegans* (*sut-2*) (Guthrie *et al*., 2009), *Sacchromyces cerevisiae* (Nab2) (Anderson *et al*., 1993; Green *et al*., 2002; Hector *et al*., 2002; Marfatia *et al*., 2003; Kelly *et al*., 2007; Kelly *et al*., 2010; Schmid *et al*., 2015; Soucek *et al*., 2016), *Sacchromyces pombe* (Nab2) (Grenier St-Sauveur *et al*., 2013), and *Drosophila melanogaster* (Nab2) (Pak *et al*., 2011; Kelly *et al*., 2016; Fasken *et al*., 2019; Lee *et al*., 2020; Corgiat *et al*., 2021; Corgiat *et al*., 2022; Rounds *et al*., 2022; Jalloh and Lancaster *et al*., 2023). These studies have collectively uncovered molecular and neuronal functions for this conserved ZnF Pab. For example, *Zc3h14* loss impairs working memory in mice where ZC3H14 protein localizes to synaptosomes in hippocampal neurons and regulates the abundance of synaptic proteins, including CaMK2α (Rha *et al*., 2017; Jones *et al*., 2020). Moreover, studies in *C. elegans* identified SUT-2 as a modulator of Tau-induced toxicity, as loss of *sut-2* robustly rescues the toxic consequences of Tau overexpression in worms (Guthrie *et al*., 2009; Currey *et al*., 2023), a function of ZC3H14/MSUT2 that extends to mice (McMillan *et al*., 2021). On the other hand, yeast *NAB2* is essential for viability (Anderson *et al*., 1993) and has critical functions in regulating transcription termination (Alpert *et al*., 2020), nuclear export (Hector *et al*., 2002), and transcript stability (Batisse *et al*., 2009; Schmid *et al*., 2015; Fasken *et al*., 2019; Alpert *et al*., 2020). Moreover, *NAB2/Nab2* loss leads to increases in bulk poly(A) tail length in yeast, mice, and flies supporting a conserved function for Nab2 in restricting poly(A) tail length (Green *et al*., 2002; Hector *et al*., 2002; Kelly *et al*., 2010; Kelly *et al*., 2014). Taken together, these findings suggest that ZC3H14/Nab2 is involved in multiple aspects of post-transcriptional RNA metabolism and that these roles may be particularly significant in neurons.

*Drosophila melanogaster* is a genetically tractable system to define molecular and developmental roles of the ZC3H14 invertebrate homolog, Nab2 (Pak *et al*., 2011). Our prior studies have determined that Nab2 function is necessary in neurons, as pan-neuronal expression of *Drosophila* Nab2 or human ZC3H14 is sufficient to rescue viability and locomotor defects associated with zygotic loss of Nab2 (Pak *et al*., 2011; Kelly *et al*., 2014). Moreover, Nab2 has a cell-autonomous role in Kenyon cells to pattern axonal projections from these cells into the mushroom bodies (Kelly *et al*., 2016), a twin neuropil structure that regulates *Drosophila* associative olfactory learning and memory (Heisenberg, 2003; Kahsai and Zars, 2011; Yagi *et al*., 2016; Takemura *et al*., 2017). Biochemical studies show that Nab2 interacts with Fmr1, the fly homolog of Fragile X Syndrome RBP, FMRP (Wan *et al*., 2000), and that these two RBPs co-regulate mushroom body morphology and olfactory memory through a mechanism likely to involve translational repression of shared Nab2-Fmr1 target RNAs (Bienkowski *et al*., 2017). Beyond the brain, Nab2 limits dendritic branching of class IV dorsal dendritic arborization (ddaC) sensory neurons through a mechanism involving the planar cell polarity (PCP) pathway (Corgiat *et al*., 2022), suggesting that Nab2 controls RNA targets encoding regulators of the actin cytoskeleton. Our recent work studying the effect of Nab2 loss on the brain transcriptome revealed that Nab2 is required for proper splicing of ~150 mRNAs (Jalloh and Lancaster *et al*., 2023). Furthermore, Nab2 limits N-6 methyladenosine (m^6^A) methylation on key mRNAs, including the alternatively spliced *Sex-lethal* (*Sxl*) transcript (Jalloh and Lancaster *et al*., 2023). However, Nab2-regulated transcripts encoding factors that guide axon and dendrite morphology have not been identified.

A recent study uncovered multiple Nab2-regulated candidate transcripts with key functions in neurodevelopment (Jalloh and Lancaster et al., 2023). Specifically, this work revealed significant retention of a 5’UTR intron in the *trio* transcript. Trio is a member of the Dbl homology (DH) family of GEF proteins with well-conserved orthologues in *C. elegans* and mammals that control F-actin polymerization through the Rac and Rho small GTPases (Bellanger *et al*., 1998b; Awasaki *et al*., 2000; Bateman *et al*., 2000; Newsome *et al.,* 2000; Bateman and Van Vactor, *2001;* Briancon-Marjollet *et al*., 2008; Ba *et al*., 2016; Pengelly *et al*., 2016; Katrancha *et al*., 2017; Backer *et al*., 2018; Bircher and Koleske, 2021). As a result of these roles, Trio loss affects axon guidance and dendritic branching as well as synaptic transmission and plasticity (Briancon-Marjollet *et al*., 2008; Iyer *et al*., 2012; DeGeer *et al*., 2015; Ba *et al*., 2016; Katrancha *et al*., 2019). Notably, *Drosophila* Trio is enriched in the brain mushroom bodies where it controls axon projection and regulates arborization of sensory ddaC neurons in the larval peripheral nervous system (PNS) (Awasaki *et al*., 2000; Iyer *et al*., 2012; Shivalkar and Giniger, 2012). Moreover, several recent studies have identified loss- and gain-of-function mutations in the human *TRIO* gene and its paralogue *KALRN* that lead to genetically dominant forms of intellectual disability and neurodevelopmental disease (Ba *et al*., 2016; Pengelly *et al*., 2016; Katrancha *et al*., 2017; Paskus *et al*., 2020).

Trio contains two GEF domains, GEF1 and GEF2, that differentially activate Rac1 or RhoA/Rho1 GTPases, respectively (Debant *et al*., 1996; Bellanger *et al*., 1998a; Bellanger *et al*., 1998b). Trio-GEF1 activation of Rac1 regulates motor neuron axon guidance, cell migration and axon outgrowth (Bateman *et al*., 2000; Newsome *et al*., 2000; Briancon-Marjollet *et al*., 2008; Peng *et al*., 2010; Song and Giniger, 2011). Comparatively, little is known about the function of Trio-GEF2; however, recent work suggests that it promotes growth cone collapse through RhoA/Rho1 (Backer *et al*., 2018). Supporting this model of opposing roles for Trio GEF1 and GEF2 function, studies in *Drosophila* ddaC neurons suggest that Trio promotes dendritic branching via GEF1 and restricts this process via GEF2 (Iyer *et al*., 2012). Despite our knowledge of Trio GEF specificity for Rac and RhoA/Rho1, how these two opposing Trio activities are modulated within axons and dendrites remains unclear.

Here, we exploit both genetic and molecular approaches to assess the role of Nab2 and the m^6^A machinery in regulating expression of the neuronally enriched protein Trio in the adult fly brain. Consistent with our previous findings that Nab2 limits m^6^A methylation on specific transcripts, reduced levels of either *Drosophila* m^6^A reader protein – the nucelar reader Yt521-B or the cytoplasmic reader Ythdf – is sufficient to rescue *Nab2^null^* viability and locomotion defects, indicating that m^6^A-mediated changes in RNA nuclear processing and cytoplasmic metabolism underlie defects in Nab2 mutants. Focusing on the *trio* mRNA, we find that Nab2 and the m^6^A methyltransferase, Mettl3, each promote an intron-excision event within the 5’UTR of a *trio* mRNA species encoding only the GEF2 (RhoGEF) domain. Intriguingly, levels of the corresponding Trio-GEF2 protein drop in heads of *Nab2^null^* but not *Mettl3^null^* flies, consistent with a model in which Nab2 modulates both nuclear splicing and cytoplasmic metabolism of the GEF2-only variant of *trio* mRNA. Critically transgenic expression of Trio-GEF2 rescues axon projection defects in *Nab2^null^* mushroom body neurons and class IV ddaC neurons while Trio-GEF1 has the opposite effect of exacerbating *Nab2^null^*neuronal defects. Together, these data identify Nab2 and Mettl3 as key regulators of *trio* 5’UTR structure and provide evidence that altered splicing and expression of Trio-GEF2 is a key driver of axon and dendrite defects in *Drosophila* lacking Nab2.

## RESULTS

### Loss of m^6^A-reader proteins rescues *Nab2^null^*defects in viability and adult locomotion

Nab2 loss causes severe defects in *Drosophila* viability, adult locomotion, and lifespan (Pak *et al*., 2011). Building on the previous finding that Nab2 loss elevates m^6^A methylation on select mRNAs (Jalloh and Lancaster *et al*., 2023), we hypothesized that some *Nab2^null^* organismal phenotypes could result from ectopic recruitment of m^6^A reader proteins onto affected mRNAs. These m^6^A reader proteins recognize m^6^A-modified adenosines via a YTH-domain (Xu *et al*., 2014) and act downstream of the methyltransferase machinery to bind and regulate the fate of methylated RNAs (Luo and Tong, 2014; Theler *et al*., 2014; Xu *et al*., 2014; Xu *et al*., 2015; Patil *et al*., 2018). Unlike more complex mammalian systems, *Drosophila* have a single nuclear m^6^A reader protein, YT-521-B (or Ythdc1) and a single cytoplasmic m^6^A reader protein, Ythdf (Haussmann *et al*., 2016; Kan *et al*., 2017).

To assess roles of nuclear Yt521-B and cytoplasmic Ythdf in *Nab2* mutant phenotypes, the *yt521-B^ΔN^* and *ythdf^ΔYTH^* alleles (Lence *et al*., 2016; Worpenberg *et al*., 2021) were individually recombined with a *Nab2^null^* allele (also known as *Nab2^ex3^*; imprecise excision of *EP3716*) (Pak *et al*., 2011) and assessed for effects on viability, adult locomotion, and lifespan. Homozygous double mutant *yt521-B^ΔN/ΔN^,Nab2^null^*flies are significantly more viable than *Nab2^null^* flies indicating that the nuclear m^6^A reader is required for the effect of Nab2 loss on viability (**Figure 1A**). *ythdf^ΔYTH^, Nab2^null^* double mutants are inviable and heterozygous reduction of cytoplasmic Ythdf (*ythdf^ΔYTH/+^, Nab2^null^*) does not improve *Nab2^null^* viability (**Figure 1A**). Reciprocally, homozygous loss of nuclear Yt521-B has no effect on *Nab2^null^*locomotion, whereas heterozygous reduction of cytoplasmic Ythdf rescues *Nab2^null^* climbing rates by approximately 6-fold as assessed in a negative geotaxis assay (at the 30s time point; **Figure 1B**). Despite the ability of reader alleles (e.g., *yt521-B^ΔN/ΔN^* or *ythdf^ΔYTH/+^*) to rescue viability or adult locomotion, neither mutant alone rescues *Nab2^null^* lifespan defects (**Figure 1C**). Together, these genetic rescue data provide evidence that neurological effects of Nab2 loss require nuclear and cytoplasmic m^6^A readers, and that each of these mechanisms may involve different mRNAs.

**Figure 1.**
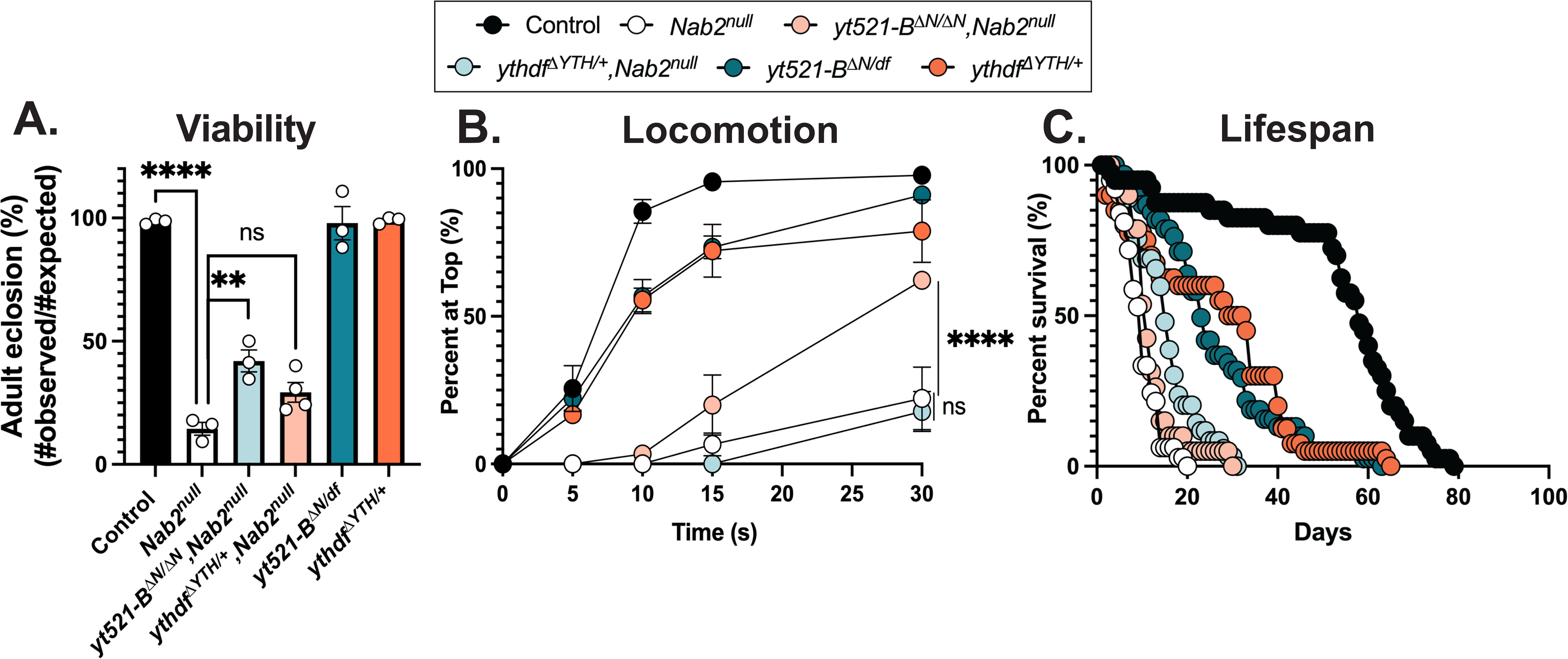
Loss of m^6^A-reader proteins rescues *Nab2^null^* defects in viability and adult locomotion. (**A**) Percent of Control, *Nab2^null^*, *yt521-B^ΔN/ΔN^Nab2^null^*, *yhdf^ΔYTH/+^Nab2^null^*, *yt521-B^ΔN/df^*, and *ythdf^ΔYTH/+^* that eclose as viable adults (calculated as #observed/#expected). (**B**) Negative geotaxis as a measure of locomotion of age-matched adult flies of indicated genotypes over time in seconds (s) taken to reach the top of a vial. (**C**) Survival of age matched adult flies of the indicated genotypes over time in days. Significance values are indicated (**p≤0.01,****p≤0.0001).

### Nab2 and Mettl3 regulate splicing of the *trio* 5’UTR in the *Drosophila* head

In light of the effects of Nab2 loss on axon and dendrite development (Kelly *et al*., 2016; Bienkowski *et al*., 2017; Rounds *et al*., 2021; Corgiat *et al*., 2022), we mined our high-throughput RNA sequencing (RNA-seq) analysis of adult heads from *Nab2^null^* mutants (imprecise excision of *EP3761*) and isogenic Controls (precise excision of *EP3716*) (Pak *et al*., 2011; Jalloh and Lancaster et al., 2023) to identify potential Nab2 target transcripts regulated by m^6^A with functions in neurodevelopment. One transcript identified in this analysis was *trio*, which encodes a Rho guanine nucleotide exchange factor (RhoGEF) that activates specific downstream Rho family GTPases (Bellanger *et al*., 1998b; Bircher and Koleske, 2021). There are multiple different variants of the *trio* transcript, two of which are readily detected in adult fly heads: hereafter referred to as *trio Medium* (*trio M*) and *trio Long* (*trio L*) (**Figure 2A, top panel**). Visualization of RNA-seq reads from *Nab2^null^* and Control heads using Integrative Genomics Viewer (IGV) (Robinson *et al*., 2017) reveals an increase in reads in introns within the 5”UTR of both *trio M* and *trio L* in *Nab2^null^* heads relative to Control (**Figure 2A**). Normal splicing patterns are detected across all other *trio* intron-exon junctions. Utilizing a publicly available me-RIP-Seq dataset from *Drosophila* heads (Kan *et al*., 2021), we bioinformatically identified three m^6^A sites in the *trio M* 5’UTR (**Figure 2A**, red lollipops), but none in the *trio L* 5’UTR. These data suggest that Nab2 is required for removal of 5’UTR introns in *trio M* and *trio L* and suggest that the removal of the *trio M* 5’UTR could also involve m^6^A.

**Figure 2.**
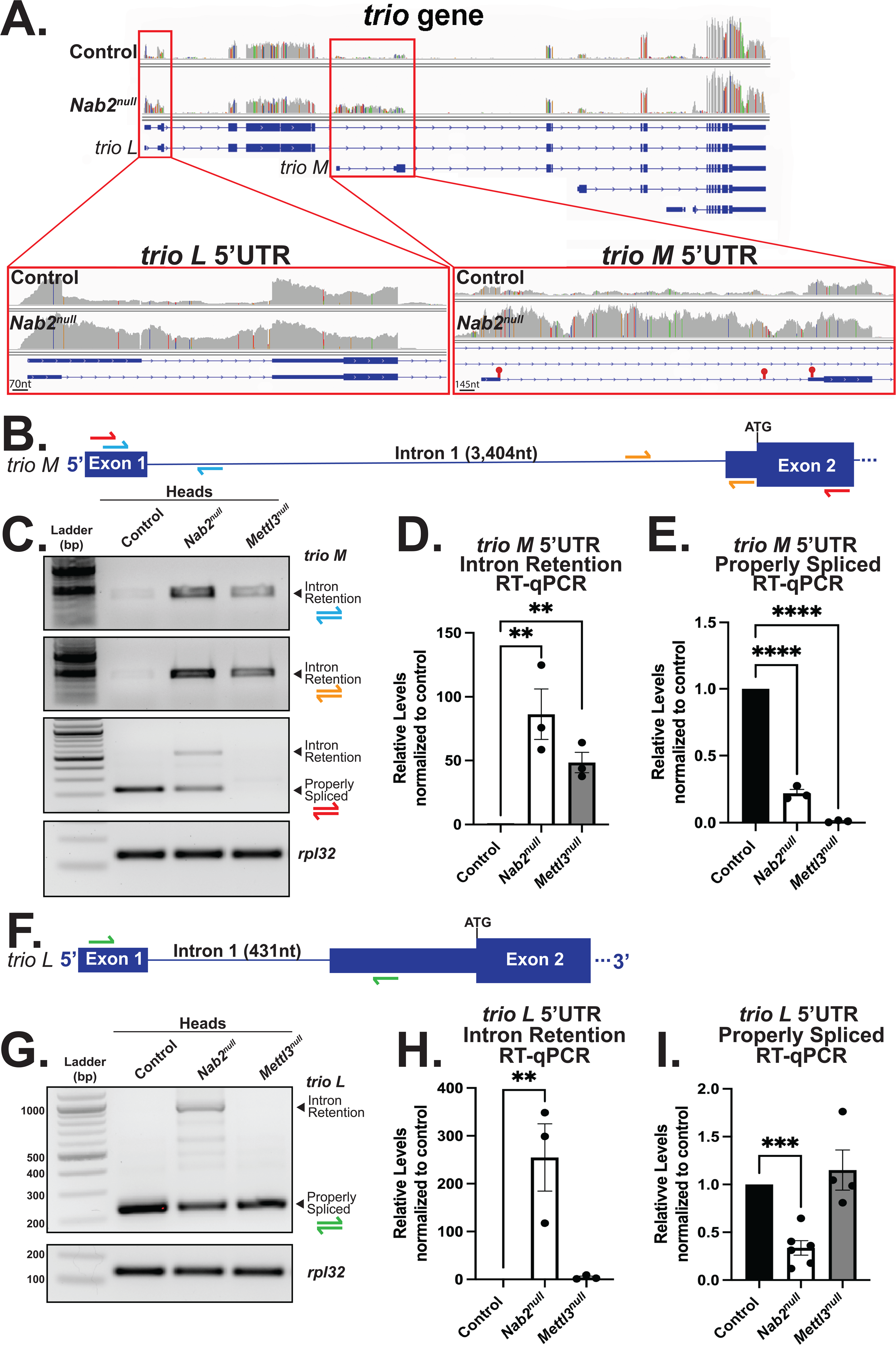
Nab2 and Mettl3 regulate splicing of the *trio* 5’UTR in the *Drosophila* head. (**A**) RNA sequencing reads across the *trio* locus in Control and *Nab2^null^* fly heads (Jalloh and Lancaster et al 2023). Boxed insets highlight the sequencing reads from the *trio L* (scale bar=70nt) and *trio M* (scale bar=145nt) 5’UTR. Red lollipops denote location of mapped m^6^A sites. (**B**) Diagram of the *trio M* 5’UTR annotated to show location of color-coded primer pairs. The position of the ATG is also indicated (**C**) RT-PCR analysis of *trio M* mRNA from Control, *Nab2^null^*, and *Mettl3^null^* heads. Properly spliced transcript and intron reattaining transcript bands are indicated. *Rpl32* serves as a control. RT-qPCR analysis detecting levels of (**D**) intron retaining or (**E**) properly spliced *trio M* transcript from Control, *Nab2^null^*, and *Mettl3^null^* heads, where control is set to 1.0. (**F**) Diagram of the *trio L* 5’UTR annotated to show location of color-coded primer pairs. (**G**) RT-PCR analysis of *trio L* mRNA from Control, *Nab2^null^*, and *Mettl3^null^* heads. Properly spliced and intron retaining transcript bands are indicated. (**H**) RT-qPCR analysis detecting levels of intron retaining and (**I**) properly spliced *trio L* transcript in Control, *Nab2^null^*, and *Mettl3^null^*heads. Significance values are indicated (**p≤0.01,***p≤0.001,****p≤0.0001).

To experimentally test this prediction, we first analyzed the *trio M* transcript using reverse transcription polymerase chain reaction (RT-PCR) analysis with primers that detect the *trio M* 5’UTR intron (exon 1-intron 1 and intron 1-exon 2) (**Figure 2B**, blue and orange primer pairs). This analysis reveals that the *trio M* 5’UTR intron-retaining transcript is enriched in adult heads of *Nab2^null^* flies as well as in heads lacking *Mettl3,* the catalytic subunit of the methyltransferase complex (**Figure 2C**) with concomitant reduction or loss of properly spliced *trio M* 5’UTR (exon 1-exon 2) (**Figure 2B**, red primer pair; see **Figure 2C**). Primers that detect correctly splied exon1-exon2 *trio M* transcript (**Figure 2B**, red primer pair) also amplify a ~550bp band in *Nab2^null^* heads that is an aberrantly spliced product corresponding to the *trio M* pre-mRNA transcript (**Figure 2B**). This RT-PCR analysis did not detect the ~4kb *trio M* 5’UTR intron-retaining transcript, possibly due to the large size of the expected product. To quantitate these results, we performed reverse transcription quantitative PCR (RT-qPCR) analysis, which confirms a significant increase in the levels of the *trio M* 5’UTR intron-retaining transcript in both *Nab2^null^* and *Mettl3^null^*heads (**Figure 2D**). Reciprocal RT-qPCR analysis to quantify the levels of properly spliced *trio M* confirms reduced levels of the properly spliced transcript in *Nab2^null^* heads and a complete loss of the properly spliced transcript in *Mettl3^null^*heads (**Figure 2E**).

Shifting the analysis to the *trio L* 5’UTR intron using primers to detect exon 1-exon 2 confirms the presence of the *trio L* 5’UTR intron-retaining transcript (exon 1-intron 1-exon 2) in *Nab2^null^* heads and reduced levels of the properly spliced transcript (exon 1-exon 2) (**Figure 2F**, green primer pair; **Figure 2G**). In contrast, only properly spliced *trio L* 5’ UTR is detected in heads of flies lacking *Mettl3* (**Figure 2G**). RT-qPCR analysis confirms increased levels of the *trio L* 5’UTR intron-retaining transcript in *Nab2^null^*, but not *Mettl3^null^*heads compared to Control (**Figure 2H**). Reciprocally, RT-qPCR analysis using primers that detect levels of properly spliced *trio L* 5’UTR show reduced transcript levels in *Nab2^null^* heads, and no change in *Mettl3^null^* heads, compared to Control (**Figure 2I**). Collectively, these data confirm that splicing of the *trio M* and *trio L* 5’UTR introns are both Nab2-dependent, but that only splicing of the *trio M* 5’UTR, and not the *trio L* 5’UTR, is Mettl3-dependent.

### Nab2 regulates levels of Trio M in the *Drosophila* head

The *Drosophila* Trio L protein is most similar to a form of human Trio protein (Trio 9S) that is enriched in the human brain and nervous system (Portales-Casamar *et al*., 2006). As illustrated in **Figure 3A**, Trio L contains a Sec14 domain, nine spectrin repeats, one Src homology 3 (SH3) domain, and two catalytic GEF domains comprised of tandem Dbl homology (DH) and pleckstrin homology (PH) domains, referred to as GEF1 and GEF2. The *Drosophila* Trio M protein corresponds to the C-terminal end of Trio L, consisting of only a SH3 domain and the GEF2 catalytic domain (**Figure 3A**).

**Figure 3.**
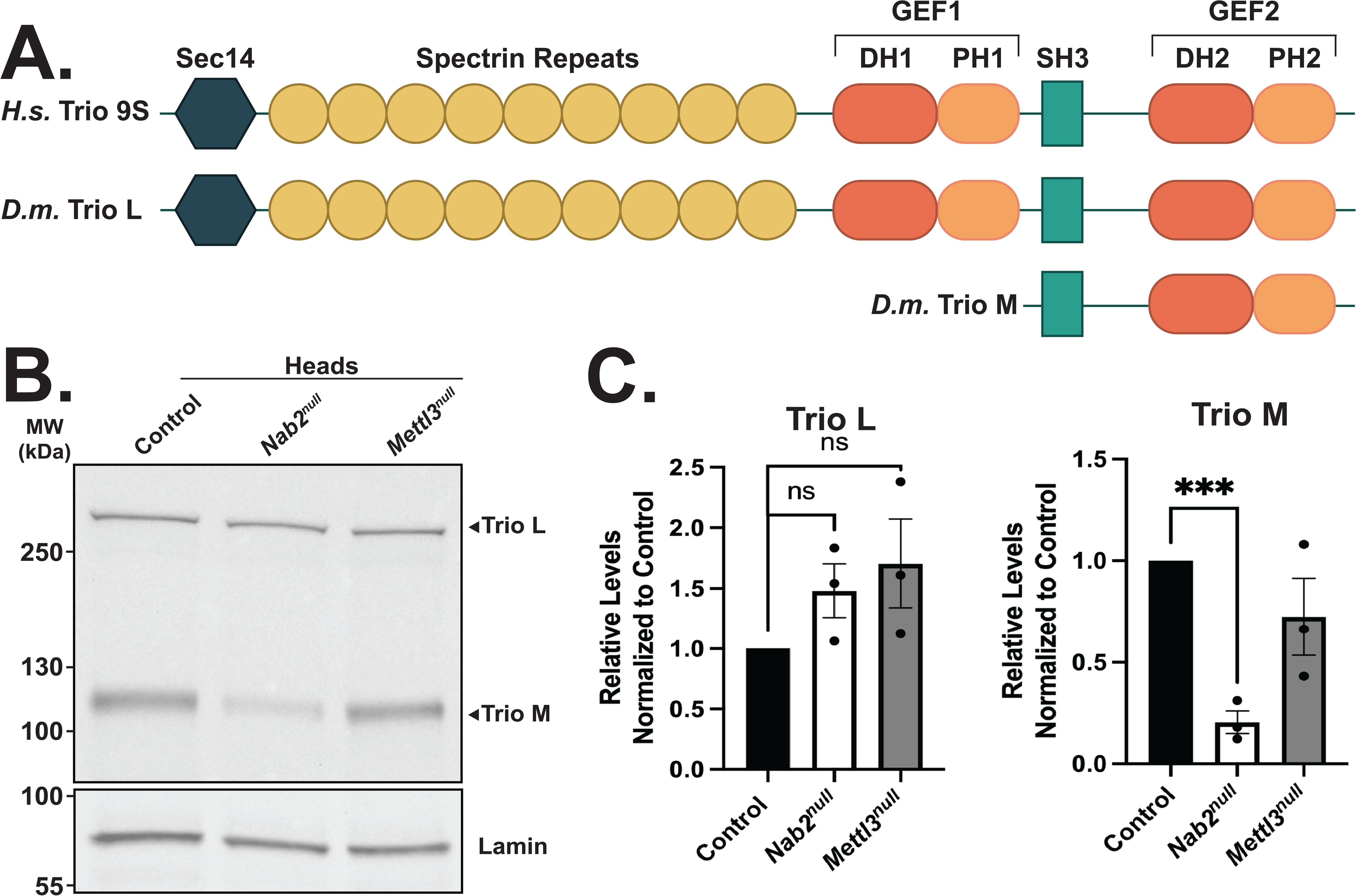
Nab2 regulates Trio M protein level in the *Drosophila* head. (**A**) Schematic of the human (*H.s.*) Trio 9S and *Drosophila melanogaster* (*D.m.*) Trio L and Trio M proteins. Trio contains a Sec14 domain, nine spectrin repeats, one Src homology 3 (SH3) domain, and two catalytic GEF domains (GEF1 and GEF2), comprised of tandem Dbl homology (DH) and pleckstrin homology (PH) domains (**B**) Immunoblotting analysis of Trio L and Trio M protein levels from Control, *Nab2^null^*, and *Mettl3^null^*heads. Lamin serves as a loading Control. Molecular weights in kDa are indicated to the left. (**C**) Quantification of Trio L (left) and Trio M (right) protein levels in (**B**) using Image Lab software. Protein levels are normalized to Control, with the value for Control set to 1.0. Asterisk denotes results that are statistically significant at **p-value≤0.001.

Based on the finding that Nab2 regulates splicing of the *trio L* and *trio M* 5’UTR introns and that Mettl3 only regulates splicing of the *trio M* 5’UTR intron, we tested whether these intron retention events affect levels of Trio L or Trio M proteins in the fly head. Immunoblotting analysis reveals that Trio L and Trio M are the major isoforms of Trio in Control brains, and that Nab2 loss reduces levels of Trio M but has no apparent effect on Trio L protein levels relative to Control (**Figure 3B**). Densitometry analysis of the Trio L and Trio M protein levels demonstrates that the decrease in the steady-state level of Trio M protein is statistically significant (**Figure 3C**). Although loss of *Mettl3* or *Nab2* results in *trio M* 5’UTR intron retention (**Figure 2C-E**), only loss of Nab2 causes a drop in Trio M protein levels (**Figure 3B&C**), implying an independent effect of Nab2 on post-splicing metabolism of the *Trio M* 5’UTR intron retaining RNA.

### Trio is altered in the *Nab2^null^* mushroom body

Previous studies demonstrated that Trio is enriched in the mushroom bodies (Awasaki *et al*., 2000), which are divided into five lobes per hemisphere (α/α’, β/β’ and γ) that project anteriorly from the dorsally located Kenyon cells (**Figure 4A**) (Heisenberg, 1998; Roman and Davis, 2001). The α/α’ lobes project dorsally, while the β/β’ and γ lobes project medially, towards the central ellipsoid body (EB) (**Figure 4A**). As demonstrated previously, loss of Nab2 causes defects in α and β lobe structures, specifically loss or thinning of the α lobes and midline fusion of the β lobe structures (Kelly *et al*., 2016; Corgiat *et al*., 2022; Rounds *et al*., 2022) (**Figure 4B**). To visualize Trio in these structures, we stained Control brains overexpressing membrane tethered GFP in α/β/γ lobes (*201Y-Gal4, UAS-mcd8::GFP*) with an α-Trio antibody, which recognizes both Trio L and Trio M protein (Awasaki *et al*., 2000) (**Figure 4B**). This analysis confirms that Trio is enriched in γ lobes, Kenyon cell bodies, and calyx as shown by colocalization with GFP, but absent or below the level of detection in α and β lobes (Awasaki *et al*., 2000) (**Figure 4B** and **Supplemental Figure 1**). Intriguingly, Trio accumulates in dysplasic axons near the midline that are not labeled with the *201Y-Gal4* driver, suggesting they are β’ lobe axons (**Figure 4B**, middle panel, bottom row). Moreover, Trio is lost in the GFP-positive γ lobes of *Nab2^null^* brains (**Figure 4B**, bottom row). Given the strong reduction of Trio M protein detected by immunoblotting of *Nab2^null^* heads (see **Figure 2B**), this result suggests that Trio M may be the primary isoform of Trio present in mushroom body γ lobes.

**Figure 4.**
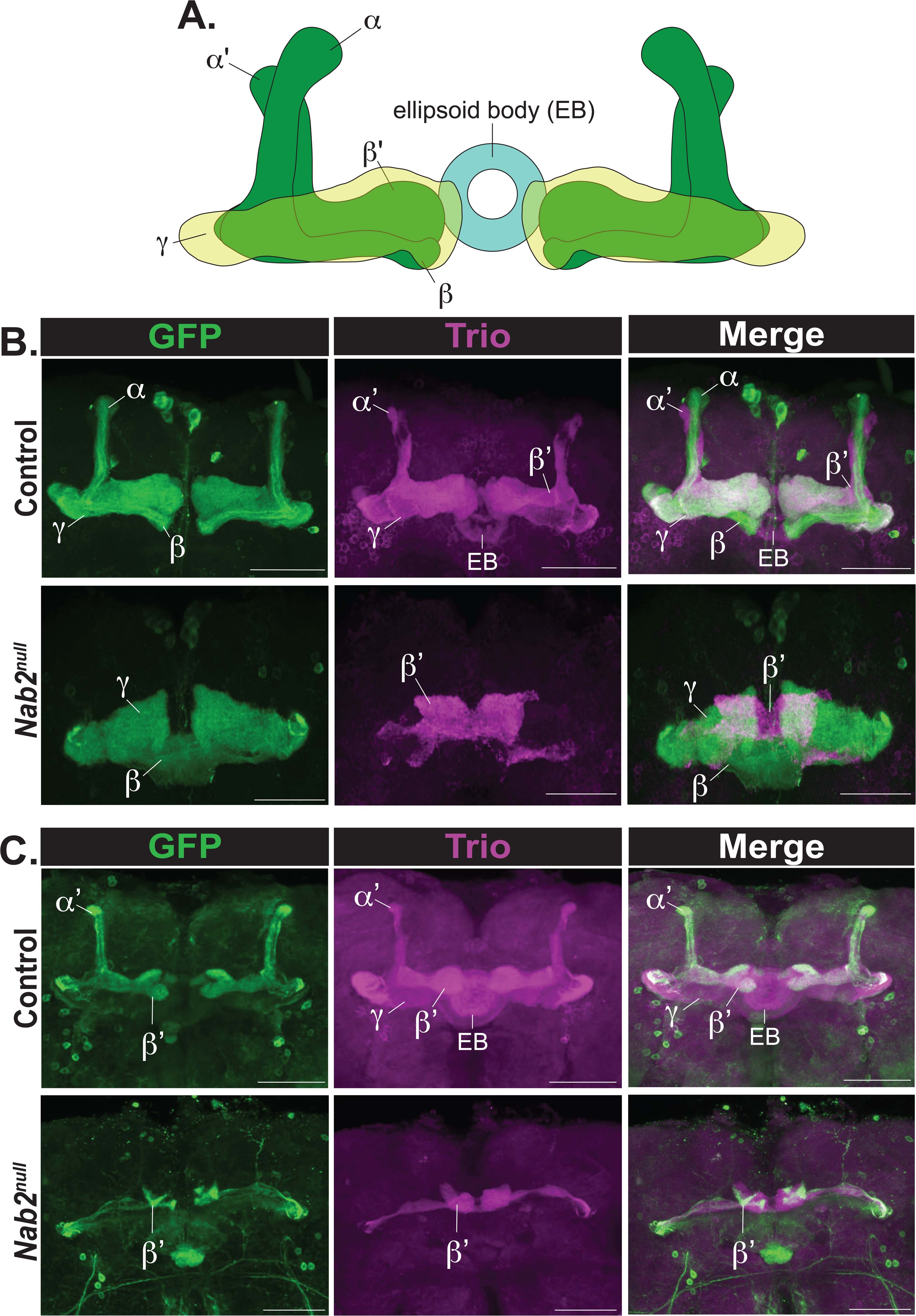
Trio is altered in the *Nab2^null^* mushroom body. (**A**) Diagram of the adult *Drosophila* mushroom body lobes depicting axons of the medially projecting gamma (γ) neurons, the vertical alpha (α) and alpha prime (α’) neurons, the medially projecting beta (β) and beta prime (β’) neurons, and the ellipsoid body (EB).(**B**) Immunofluorescence images of Control (*w-;UAS-mcd8::GFP/201Y-Gal4;;*) and *Nab2^null^* (*w-;UAS-mcd8::GFP/201Y-Gal4;Nab2^null^;*) mushroom bodies driving *UAS-mcd8::GFP* under the α, β, γ lobe-specific mushroom body *201Y-Gal4* driver. (**C**) Immunofluorescence images of Control (*w-;UAS-mcd8::GFP/C305a-Gal4;;*) and *Nab2^null^* (*w-;UAS-mcd8::GFP/C305a-Gal4;Nab2^null^;*) mushroom bodies driving *UAS-mcd8::GFP* under the α’ and β’ lobe-specific mushroom body *C305a-Gal4* driver. False colored panels show fluorescence corresponding to α-GFP (green, mcd8::GFP), α-trio (purple), and merges of the channels. Scale bar = 50 μm.

To explore the localization of Trio specifically in the α’ and β’ lobe structures, we utilized the Gal4-UAS system to overexpress GFP (*UAS-mcd8::GFP*) using a prime lobe specific Gal4 driver (*Cka^C305a^-Gal4*) (**Figure 4C**). As reported previously, Trio is enriched in α’ and β’ axons, and is also detected in γ lobe axons and ellipsoid body of Control brains (Awasaki *et al*., 2000) (**Figure 4C**, top row). *Nab2^null^* brains show complete loss or thinning of the α’ lobe axons as well as a distinct defasciculation phenotype in the β’ lobe structures (**Figure 4C**). These morphological phenotypes are accompanied by Trio loss in *Nab2^null^* γ lobe axons and ellipsoid body, and Trio accumulation in the distal portion of β’ lobe axons closest to the brain midline (**Figure 4C**, middle panel, bottom row).

### Expression of Trio GEF2 rescues α/α’ and β’ defects in *Nab2^null^* mushroom bodies

The reduced level of Trio M protein in *Nab2^null^* heads (**Figure 3B**) raises the possibility that an imbalance in the relative dose of Trio-GEF1 and Trio-GEF2 activities underlies mushroom body morphology defects observed in *Nab2^null^* flies (Kelly *et al*., 2016). This model is based on the established role of Trio protein in patterning of axons in the mushroom body (Awasaki *et al*., 2000), and predicts that loss of Trio M lowers Trio-GEF2 activity within specific mushroom body lobes.

To test this model, transgenes encoding Trio GEF1 or GEF2 domains (*UAS-Trio-GEF1* or *UAS-Trio-GEF2*) were expressed in all mushroom body lobes of *Nab2^null^* brains using the *OK107-Gal4* driver. As shown in **Figures 5A-C**, *Nab2^null^* mutant brains have highly penetrant defects in the structure of the mushroom body α lobes (missing or thinned) and β lobes (missing, thinned, or midline crossed) as detected by α-FasII staining, which specifically recognizes the α, β and weakly the γ lobes (Crittenden *et al*., 1998). Significantly, while transgenic expression of Trio-GEF2 alone has no effect on mushroom body structure in a control background this expression significantly suppresses *Nab2^null^*α lobe defects (**Figure 5A-C**). In contrast, mushroom body expression of Trio-GEF1 in control brains causes complete loss of axonal projection from the Kenyon cells (no lobe structures were detected in any of the brains analyzed; n=25) and is fully lethal in a *Nab2^null^* background (**Figure 5A-C**). Thus, Nab2 loss sensitizes mushroom body axons to the dose of Trio GEF domains such that expression of Trio-GEF2 rescues α lobe axons and addition of extra Trio-GEF1 is lethal to the animal; neither effect is observed in Control brains, indicative of a tight link between Nab2 and Trio GEF dosage in the developing mushroom body.

**Figure 5.**
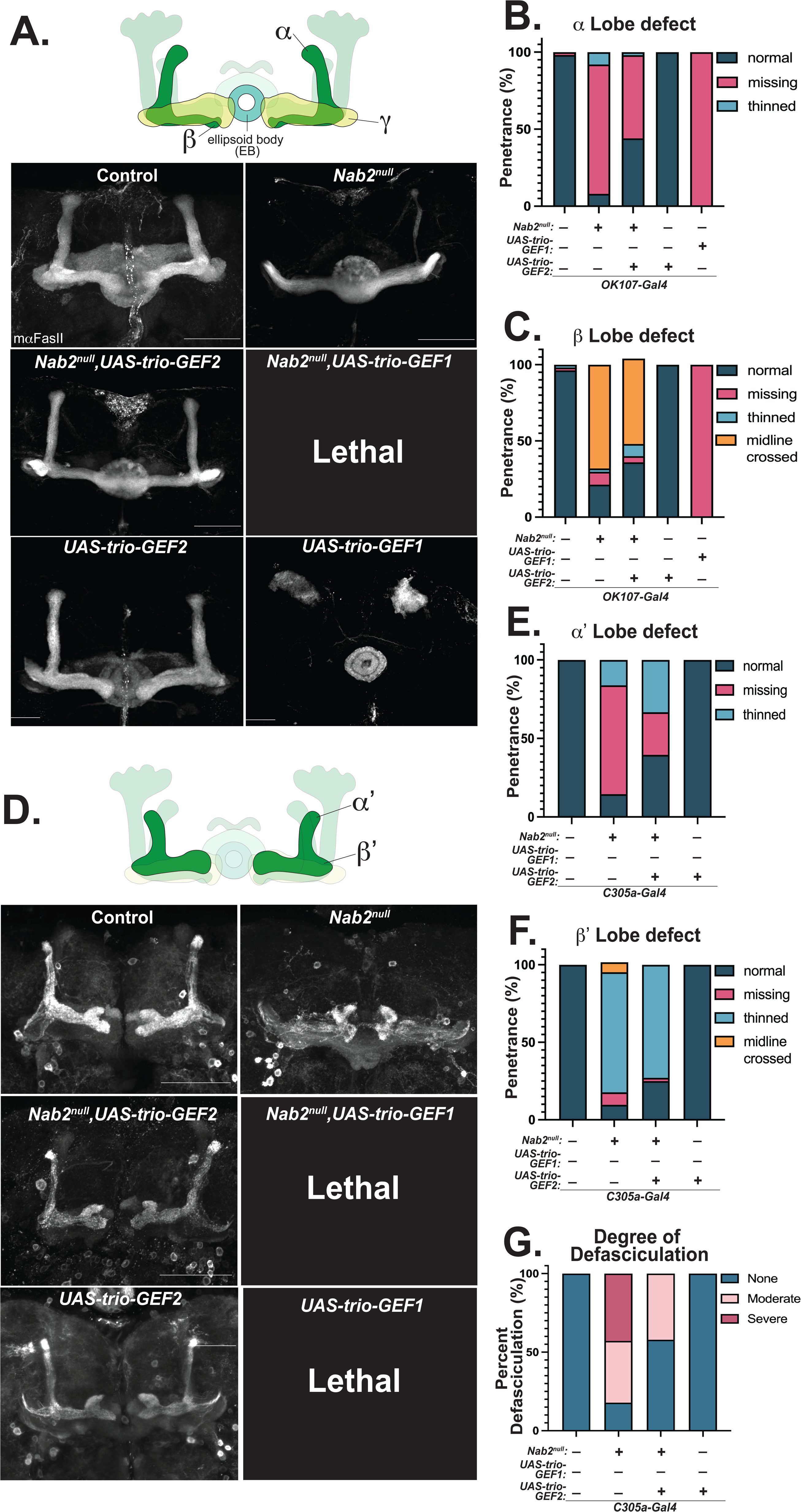
Expression of Trio GEF2 rescues α/α’ and β’ defects in *Nab2^null^* mushroom bodies. **(A**) Top: Schematic of the *Drosophila* mushroom body with lobes stained by α-Fasciclin II (FasII) highlighted. Bottom: Representative max projections of mushroom bodies of indicated genotypes stained by α-FasII. (**B**) Quantification of frequency of α lobe defects in each indicated genotype. (**C**) Quantification of frequency of β lobe defects in indicated genotypes. (**D**) Top: Schematic of the *Drosophila* mushroom body with lobes overexpressing *mcd8::GFP* under the *C305a-Gal4* driver highlighted. Bottom: Representative max projections of mushroom bodies of indicated genotypes stained with α-GFP. (**E**) Quantification of frequency of α’ lobe defects in each indicated genotype. (**F**) Quantification of frequency of β’ lobe defects in indicated genotypes. (**G**) Quantification of defasciculation phenotype severity in the indicated genotypes. Scale bar = 50 μm.

Given the enrichment of Trio protein in the α’ and β’ lobes (see **Figure 4B**), we also tested whether transgenic expression of the Trio GEF1 or GEF2 domain could autonomously rescue *Nab2^null^* α’ or β’ lobe morphology as visualized with the *Cka^C305a^-Gal4* driver (with *UAS-mcd8::GFP*). As with Trio-GEF1 or Trio-GEF2 expression in the α, β, and γ lobes using the *201Y-Gal4* driver, expression of Trio-GEF2 alone has no effect on α’ and β’ lobe morphology in Control brains, but rescues *Nab2^null^* defects in α’ lobe structure and strongly reduces β’ defasciculation (**Figure 5D-G**). Expression of Trio-GEF1 in α’ and β’ lobes has the inverse effect of late larval lethality in both Control and *Nab2^null^* animals (**Figure 5D**). These data are consistent with a model in which Nab2 acts through Trio-GEF2 to guide axon projection and fasciculation in the α’ and β’ lobes.

### Expression of Trio GEF2 rescues *Nab2^null^* dendrite defects in class IV ddaC sensory neurons

Nab2 normally restricts branching of sensory dendrites in larval class IV ddaC neurons in body wall neurons (Corgiat *et al*., 2022). Significantly, Trio-GEF1 promotes and Trio-GEF2 restricts branching of dendrites from these same class IV ddaC neurons (Iyer *et al*., 2012). Thus, we reasoned that an imbalance of Trio-GEF1 and Trio-GEF2 activities in *Nab2^null^* ddaC neurons might contribute to dendrite defects. To test this model we used a *pickpocket* (*ppk*)*-Gal4*,*UAS-mcd8::GFP* system to visualize class IV ddaC cell bodies and dendritic trees. As observed previously, loss of Nab2 (Corgiat *et al*., 2022) or transgenic expression of Trio-GEF1 (Iyer *et al*., 2012) individually increase dendritic branch complexity, while transgenic expression of Trio-GEF2 reduces dendritic branch complexity (Iyer *et al*., 2012) (**Figure 6A,B**). Significantly, combining Nab2 loss (*Nab2^null^*) with ddaC-specific expression of Trio-GEF2 rescues over-arborization normally observed in *Nab2^null^* larvae (**Figure 6A,B**). Together, these data indicate that loss of Trio-GEF2 in *Nab2^null^* larvae likely contributes to *Nab2^null^* ddaC overarborization defects.

**Figure 6.**
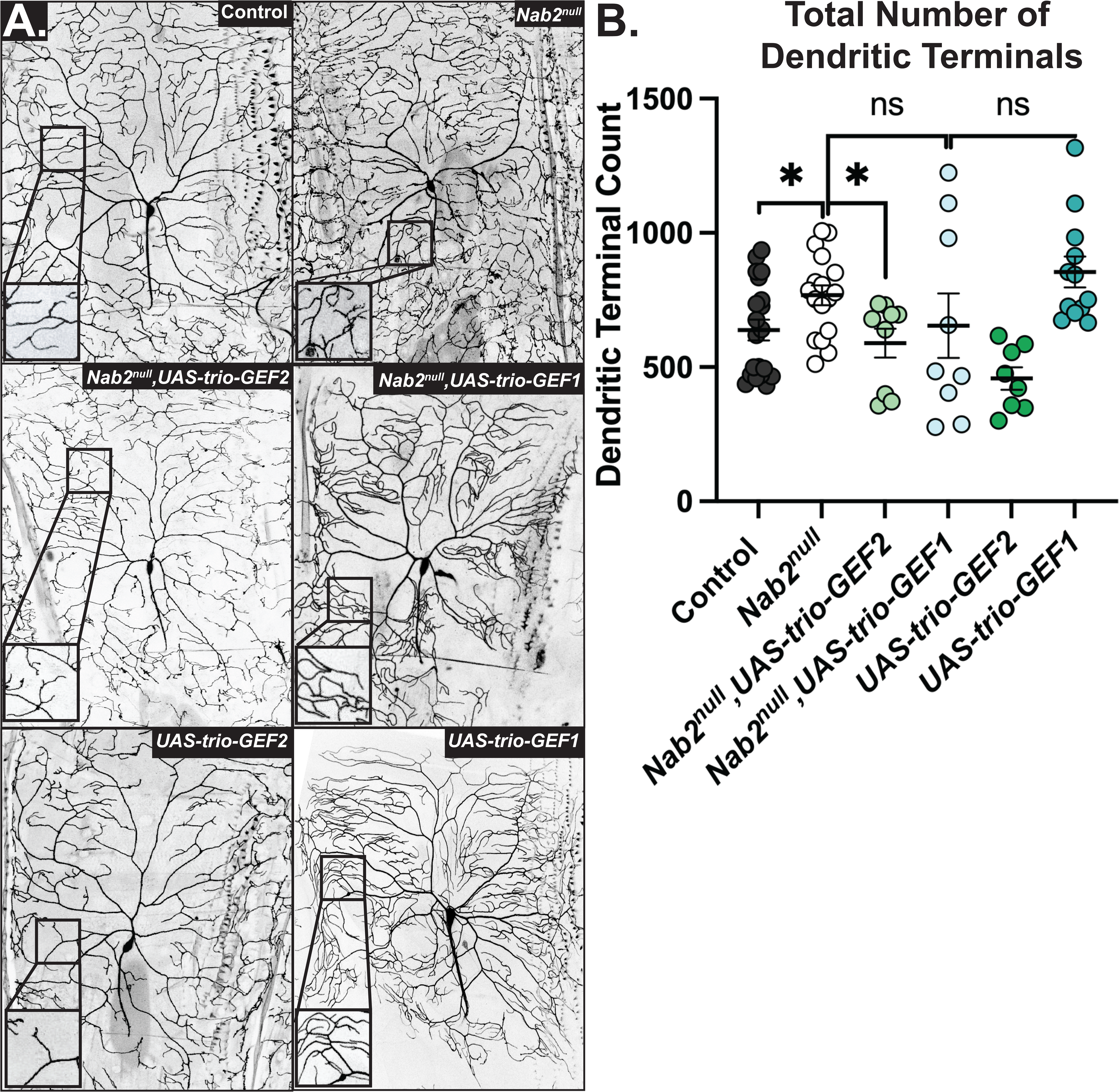
Expression of Trio GEF2 rescues *Nab2^null^* dendrite defects in class IV ddaC sensory neurons. (**A**) Maximum intensity projections of *Drosophila* class IV ddaC neurons from Control, *Nab2^null^, Nab2^null^ UAS-Trio-GEF2, Nab2^null^ UAS-Trio-GEF1, UAS-Trio GEF2, and UAS-Trio-GEF1* L3 larvae. Inset black boxes show high magnification views of dendritic arbors. (**B**) Quantification of total number of dendritic terminals for each genotype. Asterisk denotes results that are statistically significant at *p-value≤0.05.

### Expression of Trio GEF2 rescues *Nab2^null^* defects in viability and locomotion

Given the genetic interactions between the *UAS-Trio-GEF1* and *UAS-Trio-GEF2* transgenes and *Nab2*, we assessed whether transgenic expression of these distinct Trio GEF domains in the mushroom body (*OK107-Gal4, UAS-Trio-GEF1* or *UAS-Trio-GEF2*) affects the organismal phenotypes of viability, adult locomotion, and lifespan. Confirming our previous findings (Pak *et al*., 2011), we detected severe reductions in viability, adult locomotion, and lifespan in *Nab2^null^* flies compared to Control (**Figure 7A-C**). As previously noted, expression of Trio-GEF1 in mushroom bodies is lethal at late larval stages (**Figure 7A**). Remarkably, transgenic expression of Trio-GEF2 in mushroom bodies strongly suppresses *Nab2^null^* defects in viability and suppresses defects in adult locomotion but does not significantly alter lifespan (**Figure 7A-C**), indicating that a deficit in Trio-GEF2 regulated cytoskeletal dynamics within *OK107*-expressing brain neurons contributes to developmental and post-developmental defects in *Drosophila* lacking Nab2.

**Figure 7.**
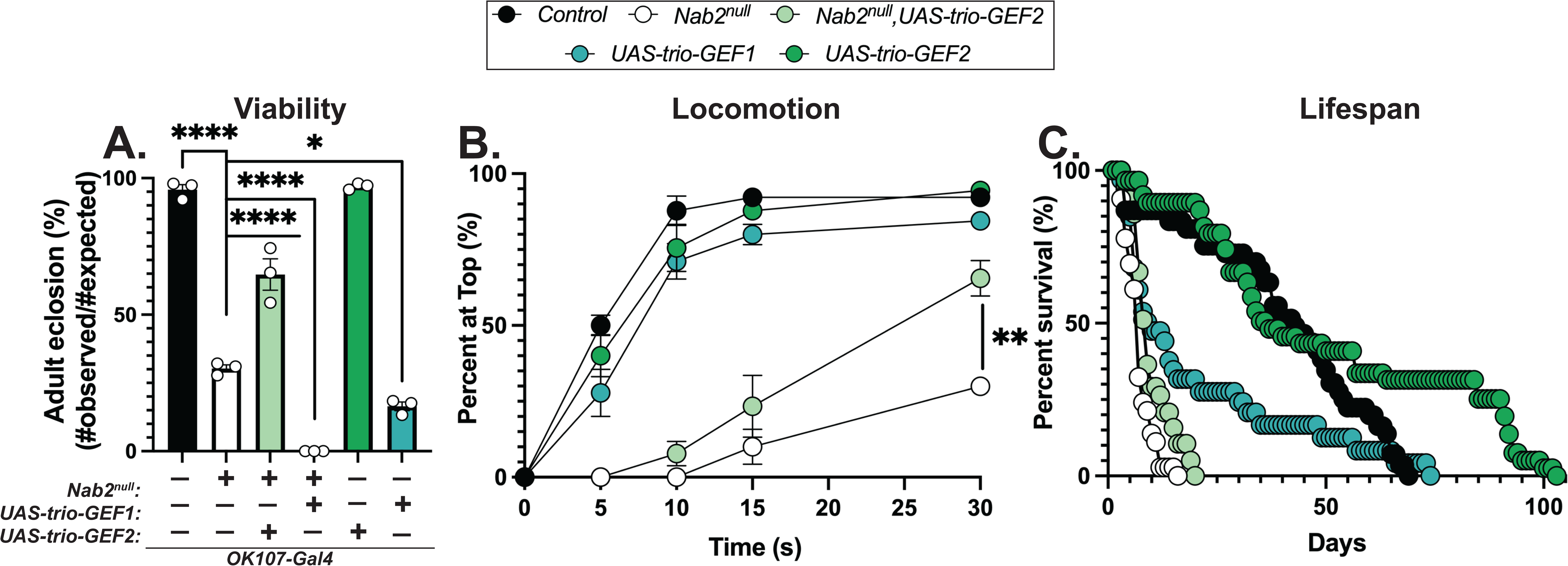
Expression of Trio GEF2 rescues *Nab2^null^* defects in viability and locomotion. (**A**) Percent of Control, *Nab2^null^*, *Nab2^null^ UAS-Trio-GEF2, UAS-Trio GEF2,* or *UAS-Trio-GEF1* that eclose as viable adults (calculated as #observed/#expected) using the *OK107-Gal4* mushroom body driver. (**B**) Negative geotaxis of age-matched adult flies of indicated genotypes over time in seconds (s). (**C**) Survival of age matched adult flies of the indicated genotypes over time in days. Significance values are indicated (*p≤0.05,**p≤0.01,****p≤0.0001).

## DISCUSSION

Here, we identify a role for the *Drosophila* Nab2 RBP and Mettl3 m^6^A methyltransferase in regulating the *trio* mRNA, which encodes a conserved RhoGEF protein that is mutated in human intellectual disability and regulates axon guidance and dendritic arborization through two GEF domains that individually control the cytoskeletal regulators Rac and RhoA/Rho1 (Debant *et al*., 1996; Bellanger *et al*., 1998a; Bellanger *et al*., 1998b; Bateman and Van Vactor, 2001; Jaffe and Hall, 2005; Briancon-Marjollet *et al*., 2008; Iyer *et al*., 2012; Bircher and Koleske, 2021). The data presented provide strong evidence that the *trio* transcript is a key downstream target of Nab2 in neurons based on an m^6^A- and Nab2-dependent splicing event and identifies specific effects of each Trio GEF domain within axons and dendrites that develop from neurons lacking Nab2. The results of this study combine with our previous work (Corgiat *et al*., 2022; Jalloh and Lancaster *et al*., 2023) to support a model in which Nab2 regulates transcripts that encode key regulators of neurodevelopment, including the conserved GEF Trio. In the broader context, the phenotypic consequences of loss of an RBP result from the collective changes to target transcripts, and defining the mechanistic basis of these phenotypes requires systematic analysis of how individual targets are impacted. In the case of Nab2, evidence now supports both m^6^A-dependent and independent roles in *trio* mRNA splicing as well as potential effects on cytoplasmic metabolism of *trio* mRNA. Taken together, these findings support a model where RBPs such as Nab2/ZC3H14 regulate a collection of target transcripts, potentially through multiple mechanisms, that all contribute to downstream phenotypes.

Previous work illustrating broad rescue of *Nab2^null^*phenotypes by Mettl3 heterozygosity (Jalloh and Lancaster *et al*., 2023) suggested that other regulators of m^6^A-modified transcripts could also contribute to *Nab2^null^* defects in viability, adult locomotion, and lifespan. Here, we demonstrate that loss of nuclear or cytoplasmic m^6^A reader function rescues some, but not all, organismal phenotypes associated with loss of Nab2 (**Figure 1**). Homozygous loss of the nuclear m^6^A reader YT521-B suppresses *Nab2^null^* defects in viability, but not locomotion or lifespan, whereas heterozygous loss of the cytoplasmic m^6^A reader Ythdf dominantly suppresses *Nab2^null^* defects in locomotion, but not viability or lifespan. Collectively, these data suggest that mRNA targets of Nab2 responsible for these behavioral phenotypes may be differentially regulated between cell types and in an m^6^A-dependent manner. For instance, Nab2-regulated transcripts encoding proteins that govern *Drosophila* viability could rely more heavily on nuclear m^6^A regulatory mechanisms, such as splicing or export. On the other hand, Nab2-regulated transcripts encoding proteins that govern *Drosophila* negative geotaxis could more heavily require cytoplasmic m^6^A regulatory mechanisms, such as translation or stability. Moreover, the inability of *YT521-B* or *Ythdf* loss to rescue *Nab2^null^* defects in lifespan suggests that Nab2-regulated transcripts that govern lifespan may not by modified by m^6^A, or that Nab2 plays m^6^A-independent roles in regulating transcripts critical to control lifespan.

Previous studies demonstrated that loss of m^6^A regulatory proteins disrupts axon projection in the *Drosophila* mushroom body (Kan *et al*., 2021; Worpenberg *et al*., 2021). Although *Mettl3* heterozygosity broadly rescues *Nab2^null^* behavioral defects (Jalloh and Lancaster *et al*., 2023), it does not dominantly suppress mushroom body morphology defects (**Supplemental Figure 2**). This finding suggests that Mettl3 heterozygosity is insufficient to reduce m^6^A methylation on Nab2-target transcripts to a degree necessary for rescue of axonal projection defects. Given the ability of m^6^A writer and reader alleles to broadly rescue *Nab2^null^* phenotypes, future studies will aim to further define the relationship between m^6^A machinery and Nab2 in relation to regulation of *Drosophila* mushroom body morphology. Moreover, a genome-wide approach to assess transcriptomic changes in m^6^A in *Nab2^null^* flies will help delineate the neuronal, Nab2-regulated transcripts that exhibit changes in m^6^A methylation.

A number of *trio* transcript variants exist in the *Drosophila* brain (Awasaki *et al*., 2000). Here, we demonstrate that both Nab2 and Mettl3 are required for proper splicing of the 5’UTR intron of *trio M*. On the other hand, splicing of the 5’UTR intron of *trio L* is dependent on Nab2 but not Mettl3. These data align with previously published RNA-seq data from *Nab2^null^ Drosophila* heads (Jalloh and Lancaster *et al*., 2023), as well as a publicly available mi-CLIP-seq dataset that mapped m^6^A sites in *trio M* 5’UTR intron, but not *trio L* 5’UTR intron (Kan *et al*., 2021). Notably, our data suggest that disruptions in splicing of the *trio* 5’UTR by loss of Nab2 or Mettl3 do not necessarily correspond with perturbations in steady-state levels of Trio L and Trio M. These results suggest that the intron retention event in the 5’UTR of *trio M* results in a significantly reduced level of Trio M protein in the *Nab2^null^* fly head. Surprisingly, the steady-state level of Trio M protein is unaffected in *Mettl3^null^* heads even though the 5’UTR intron retention is comparable to the levels observed in *Nab2^null^* heads. Given previously defined roles for Nab2 as a potential inhibitor of m^6^A methylation (Jalloh and Lancaster *et al*., 2023), this observation suggests that excess m^6^A on the *trio M* pre-mRNA upon loss of Nab2 may disrupt subsequent translation, trafficking, or stability. In contrast, retention of the *trio L* 5’UTR intron that occurs upon loss of Nab2 does not affect the steady-state level of the Trio L protein, suggesting this intron retention event may not disrupt translation. Alternatively, the remaining level of the properly spliced *trio L* in *Nab2^null^*heads may be sufficient to maintain the steady-state level of Trio L protein.

Our results support a model where loss of Trio M, and therefore GEF2 function, contributes to morphological defects in mushroom bodies of *Nab2^null^* flies. Previous studies demonstrated that Trio is a critical regulator of mushroom body morphology and Trio is enriched in the α’, β’ and γ lobes, but is virtually absent in the α and β lobes (Awasaki *et al*., 2000). We confirm these findings and further demonstrate that upon loss of Nab2, Trio levels are depleted in the γ lobes. Despite the established functions of Trio in regulating γ lobe formation (Awasaki *et al*., 2000), γ lobe defects are not detected upon loss of Nab2 (Kelly *et al*., 2016). Given that Trio M is the only isoform of Trio depleted in *Nab2^null^*brains (**Figure 3B**) and γ lobes show no defects (**Figure 5A**), this finding suggests that Trio L, and therefore GEF1 function, is responsible for patterning γ lobe axons in the developing brain.

Studies of axon pathfinding mechanisms in the mushroom body demonstrate that the α’ and β’ lobes guide development of the α and β lobes (Fushima and Tsujimura, 2007). Given that Trio is enriched in the α’/β’/γ lobes of the mushroom body, our data suggest that loss of Trio M, and therefore GEF2 levels, in α’/β’ lobes may contribute to *Nab2^null^* α/β lobe defects. We demonstrate that transgenic expression of Trio-GEF2 in α’/β’ lobes (*C305a-Gal4*) of *Nab2^null^* mushroom bodies rescues α’ lobe defects and β’ lobe defasciculation phenotypes in a cell autonomous manner (**Figure 5**). Moreover, expression of Trio-GEF2 in all mushroom body lobes (*OK107-Gal4*) rescues *Nab2^null^* α lobe defects; however, whether this rescue occurs in a cell autonomous manner remains unknown (**Figure 5**). Collectively, these data demonstrate that Trio-GEF2 rescue is limited to some mushroom body lobes (α, α’, β’) and not others (β), implying that Trio-GEF2 is required for projection of only some *Nab2^null^* axons. In this regard, misprojection defects in *Nab2^null^* β axons are not rescued by multiple genetic manipulations that rescue α axon defects (e.g., by transgenic expression of Trio-GEF2 [this study] or by single copy alleles of *fmr1*, *Atx2*, or PCP components (Kelly *et al*., 2016; Bienkowski *et al*., 2017; Rounds *et al*., 2021; Corgiat *et al*., 2022)). These findings imply roles for Nab2 in these two types of Kenyon cell projections, and suggest that Nab2 regulates different mRNAs to govern development of distinct mushroom body lobes.

Previous studies have established that the Trio-GEF1 domain acts primarily through activation of Rac1 to promote axon outgrowth and pathfinding, while Trio-GEF2 acts primarily through RhoA/Rho1 to restrict neurite outgrowth (Leeuwen *et al*., 1997; Iyer *et al*., 2012; Bircher and Koleske, 2021). Given that loss of Nab2 disrupts the ratio of GEF1 and GEF2 in *Drosophila* heads by decreasing the level of Trio M but not Trio L, we hypothesized that expression of the GEF1 effector, Rac1, in mushroom body neurons would exacerbate *Nab2^null^* phenotypic and morphological defects, whereas expression of the GEF2 effector, RhoA/Rho1, would rescue these same defects. Intriguingly, we observed that expression of either Rac1 or RhoA/Rho1 in the absence of Nab2 in Trio-enriched mushroom body neurons is lethal (**Supplemental Figure 3**). These data suggest that further expression of Rac1 in *Nab2^null^*flies in which GEF1 levels, and therefore likely Rac1 activation, dominates is detrimental to nervous system development. On the other hand, the lethality induced by expression of RhoA/Rho1 upon loss of Nab2, indicates that Trio-GEF2 may act via other unknown effectors to govern mushroom body development.

Nab2 and Trio have established roles in sculpting dendritic arborization of class IV ddaC neurons in the *Drosophila* peripheral nervous system (Iyer *et al*., 2012; Corgiat *et al*., 2022). Here, we demonstrate that transgenic expression of the Trio-GEF2 domain rescues overarborization defects in *Nab2^null^*class IV ddaC neurons. In line with previous studies (Iyer *et al*., 2012), we validate that transgenic expression of Trio-GEF1 in class IV ddaC neurons causes dramatic overarborization defects, while transgenic expression of Trio-GEF2 causes underarborization defects compared to control animals. Interestingly, we also demonstrate that expression of Trio-GEF1 in class IV ddaC neurons of *Nab2^null^* flies results in a wide range of arborization phenotypes. Very few of these animals survive to the wandering 3^rd^ instar larval stage and no animals survive to adulthood. Given these observations, disruption of Nab2-regulated mRNAs in these neurons as well as over-activation of Rac1 by GEF1 may severely disrupt ddaC development such that arborization defects are highly variable from animal-to-animal. Overall, these data support a role for Trio M, and therefore GEF2 loss, in contributing to the established overarborization defects in *Nab2^null^* class IV ddaC neurons (Corgiat *et al*., 2022).

In aggregate, these data reveal a role for Nab2 and Mettl3 in regulating splicing and protein levels of the RhoGEF Trio to support proper nervous system development. Genetic interactions between the m^6^A machinery and Nab2 support a role for Nab2 in the regulation of m^6^A methylation. We show for the first time that loss of Trio M, and therefore GEF2 levels, in *Nab2^null^* flies contributes to several *Nab2^null^*defects, including neuronal defects, such as mushroom body morphology, class IV ddaC arborization. Moreover, we demonstrate that transgenic expression of Trio-GEF2 broadly rescues *Nab2^null^* viability and adult locomotion. This regulatory relationship between Nab2 and Trio-GEF2 could be cell autonomous and may also suggest interactions between neurons and their surrounding environment. Given that mutations in human *ZC3H14* and *TRIO* are both linked to intellectual disabilities (Pak *et al*., 2011; Ba *et al*., 2016), dysregulation of Trio function in neurons is one potential mechanism to explain axonal and dendritic phenotypes observed in *Nab2^null^ Drosophila* (Pak *et al*., 2011; Kelly *et al*., 2016; Bienkowski *et al*., 2017; Corgiat *et al*., 2021; Corgiat *et al*., 2022; Jalloh and Lancaster *et al*., 2023) and *Zc3h14* mutant mice (Jones *et al*., 2020), as well as the cognitive defect observed in human patients lacking ZC3H14 (Pak *et al*., 2011).

## METHODS

### RESOURCES TABLE

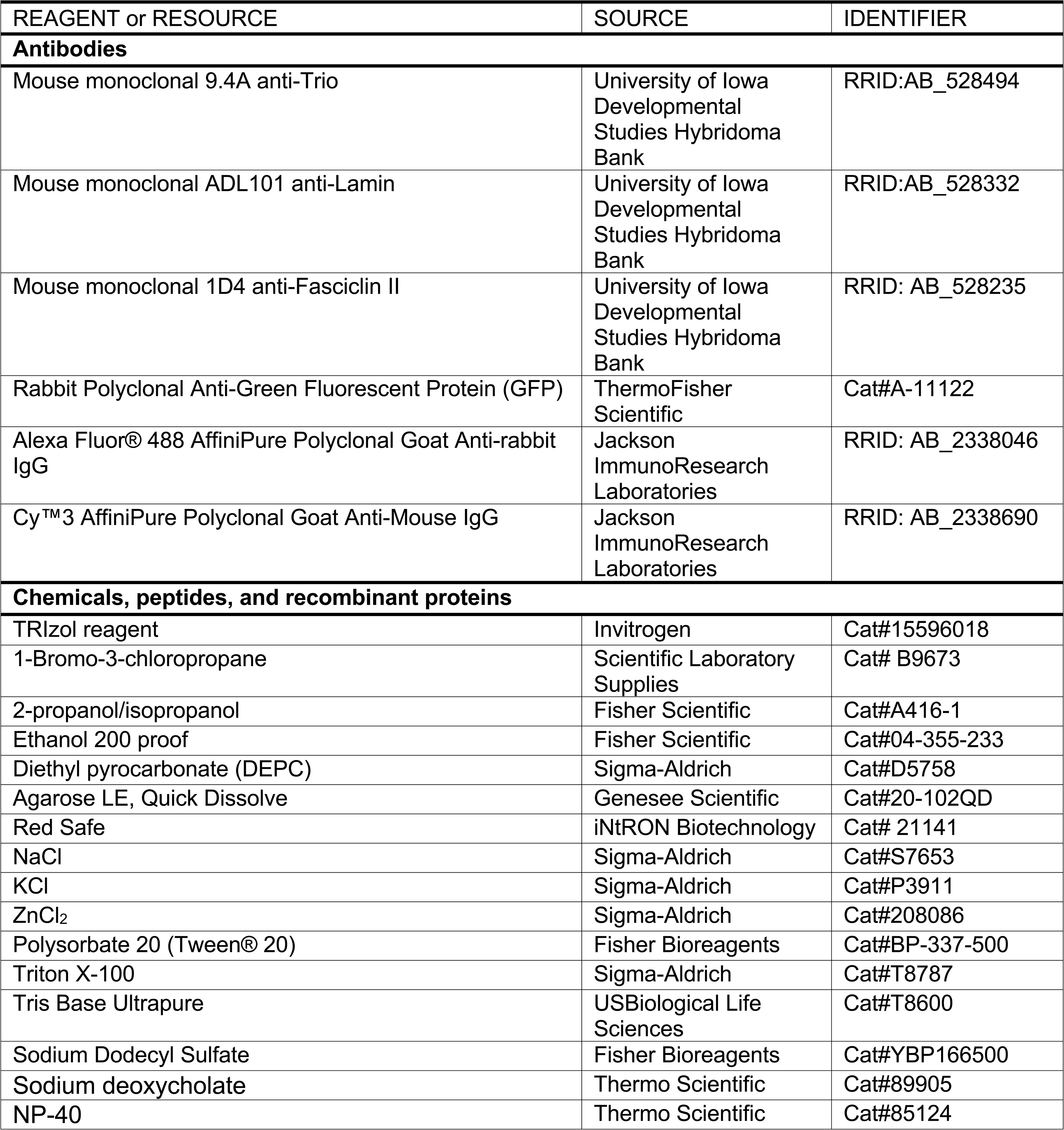
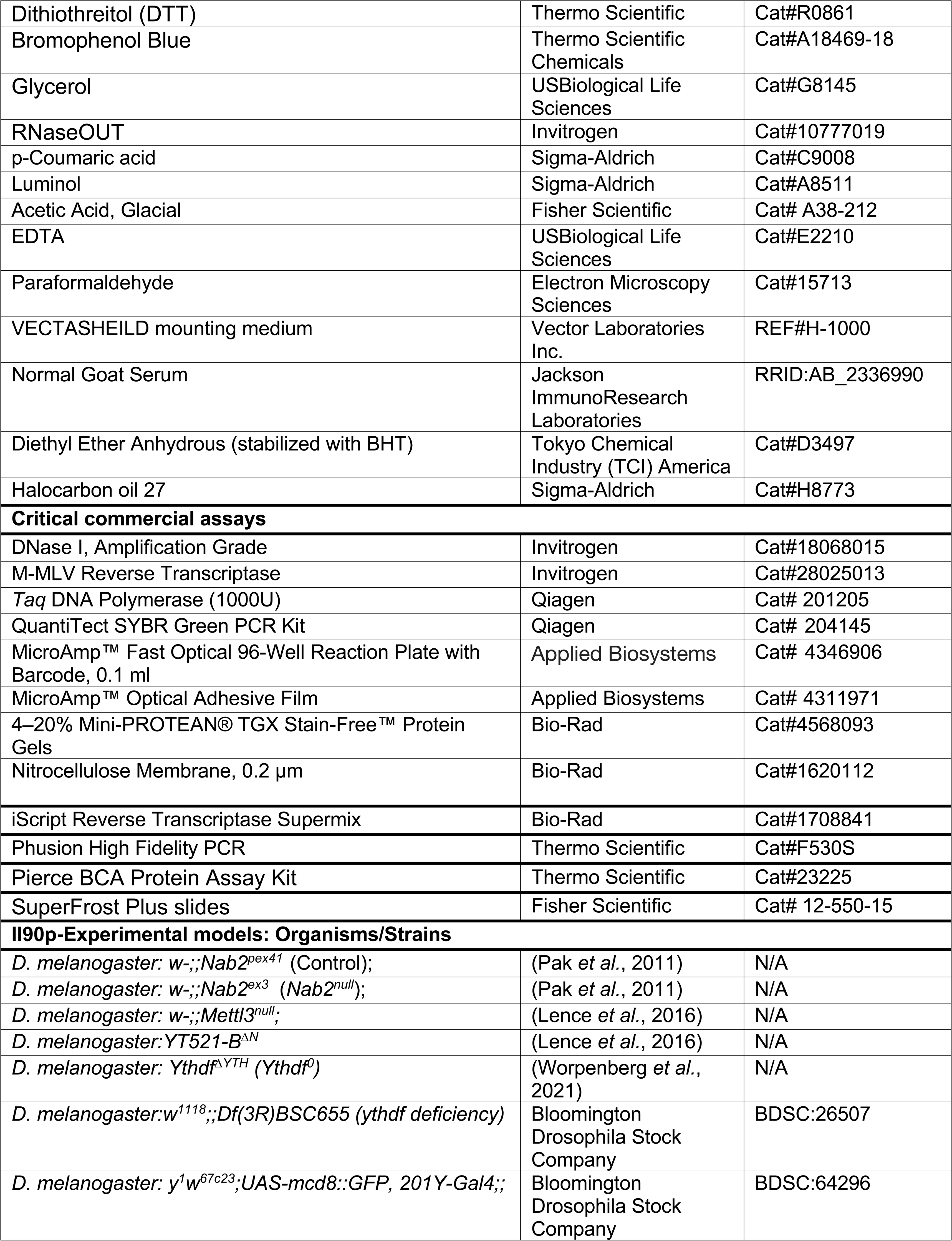
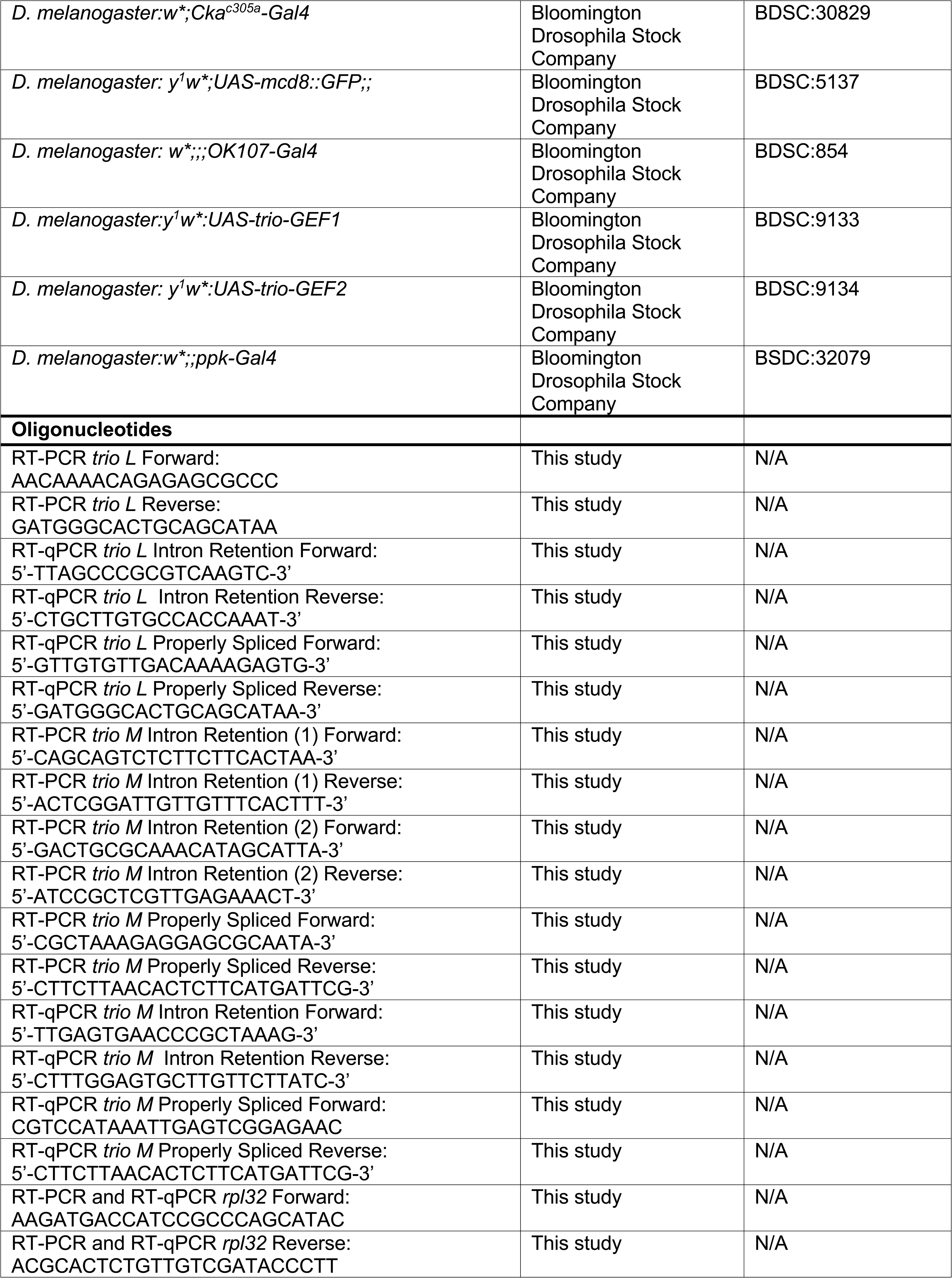
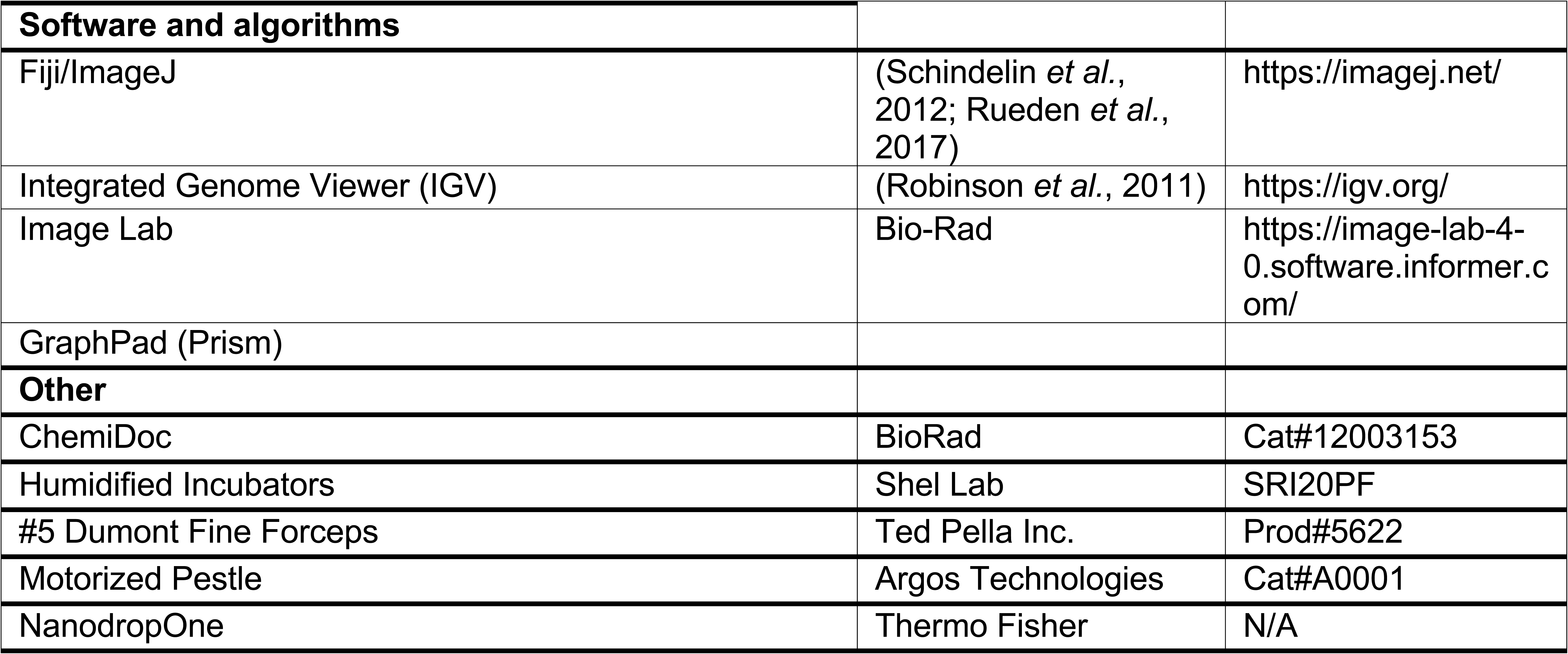

### RESOURCE AVAILIABILTY

#### Lead contact

Further information and requests for resources and reagents should be directed to and will be fulfilled by the Lead Contact, Ken Moberg.

#### Materials availability

The *Drosophila melanogaster* lines generated in this study are available by contacting the Lead Contact.

#### Data and code availability

This study did not generate any dataset or codes.

### EXPERIMENTAL MODEL AND SUBJECT DETAILS

Flies (*Drosophila melanogaster)* were raised on standard cornmeal agar medium and maintained in an incubator set at 25°C with a 12-hour light/dark cycle. Crosses were reared under the same conditions, and standard medium was supplemented with dry yeast. The GAL4-UAS binary transgenic system was used to express transgenes of interest. Details of genotypes can be found in the Key Resources Table. One to 5 day-old flies were used for experiments in this study. An equal number of males and females was used for all experiments.

### METHOD DETAILS

#### *Drosophila melanogaster* stocks and genetics

*Drosophila melanogaster* stocks were raised on standard cornmeal agar and maintained in humidified incubators (SRI20PF, Shel Lab) at 25°C with 12-hour light/dark cycles. Crosses were reared under the same conditions and supplemented with dry yeast. The strains used in this study are described in the Key Resources Table.

#### Viability and lifespan analysis

Viability at 25°C was measured by assessing eclosion rates of 100 wandering L3 larvae collected for each genotype, and then reared in a single vial. Hatching was recorded for 5-6 days. At least three independent biological replicates per genotype were tested for significance and calculated using group analysis on GraphPad (Prism). Lifespan was assessed at 25°C as described previously (Morton *et al*., 2020). In brief, newly eclosed flies were collected, placed in vials (10 flies per vial), and then transferred to fresh vials weekly, or as needed). Survivorship was scored daily. At least three independent biological replicates were tested for each genotype, and significance was calculated using group analysis on GraphPad (Prism).

#### Locomotion assays

Negative geotaxis was tested as previously described (Morton *et al*., 2020). Briefly, newly eclosed flies (day 0) were collected, and kept in vials for 2-5 days. Cohorts of 10 age-matched flies were transferred to a 25 ml graduated cylinder for analysis. Flies in graduated cylinders were tapped to bring flies to the bottom of the vial and the rate at which the flies traveled to the top of the vial (25 ml mark) was measured at 5, 10, 15, and 30s). At least three biological replicates per genotype were analyzed and significance was calculated using grouped analysis on GraphPad (Prism).

#### *Drosophila* decapitation

CO_2_-anesthetized flies were collected and frozen at −80°C for approximately five minutes. Frozen flies were then placed on a metal plate over dry ice. Gently, #5 Dumont fine forceps (Ted Pella, Inc.) were placed between the *Drosophila* head and thorax to remove the head from the remainder of the body. Heads were carefully placed in Eppendorf tubes, on ice, for subsequent processing.

#### RNA isolation for RT-PCR and real-time qPCR

Total RNA was isolated from adult heads using the TRIzol (Invitrogen) method. Briefly, *Drosophila* heads were homogenized in 0.1 ml TRIzol using a motorized pestle (Argos Technologies) on ice. TRIzol was added to samples to bring to a total volume of 0.5 ml and 0.1 ml of 1-Bromo-3-chloropropane (Scientific Laboratory Supplies) was added. Samples were vortexed on high speed for 10s and incubated at room temperature for 15 min. Next, samples were centrifuged for 15m at 13,000 x g at 4°C. The top, aqueous layer was removed and placed into a clean Eppendorf tube. An equal volume of 2-propanol (~250 μL) was added. Samples were inverted 10 times and incubated at room temperature for 10 min. Next, samples were centrifuged for 15m at 13,000 x g at 4°C. The supernatant was removed, and 0.5 ml of 75% ethanol was added. Samples were centrifuged a final time for 15 min at 13,000 x g at 4°C. The supernatant was removed, and the samples were allowed to air dry until the remaining ethanol evaporated (~5 min). The pellet was resuspended in 10-20 μL of DEPC water. RNA concentration and purity was assessed using a Spectrophotometer (Thermo Fisher). Total RNA (1 μg) was treated with DNaseI (Qiagen). cDNA was generated using M-MLV Reverse Transcriptase (Invitrogen). Qiagen *Taq* polymerase (Qiagen) was used for PCR amplification of target transcripts and products were resolved and imaged on 1-2% agarose gels (Chemi-Doc). Quantitative real-time PCR reactions were carried out in technical triplicate with QuantiTect SYBR Green Master Mix using Applied Biosystems StepOne Plus real-time machine (ABI). Results were analyzed using ΔΔCT method, normalized to loading Control (e.g., *rpl32*), and plotted as relative levels normalized to Control. Primers used for all PCR reactions are listed in the Key Resources Table.

#### Immunoblotting

For analysis of Trio protein levels, heads of newly eclosed flies were decapitated and collected on dry ice. Protein lysates were prepared by homogenizing heads in 0.5 ml of RIPA-2 Buffer (50 mM Tris-HCl, pH 8; 150 mM NaCl; 0.5% sodium deoxycholate; 1% Igepal CA-630 0.1% SDS) supplemented with protease inhibitors (1 mM PMSF; Pierce Protease Inhibitors; Thermo Fisher Scientific) and 1% SDS. Samples were sonicated 3 x 10 s with 1 min on ice between repetitions, and then centrifuged at 13,000 x g for 15 min at 4°C. Protein lysate concentration was determined by Pierce BCA Protein Assay Kit (Life Technologies). Head lysate protein samples (40–60 μg) in reducing sample buffer (50 mM Tris-HCl, pH 6.8; 100 mM DTT; 2% SDS; 0.1% Bromophenol Blue; 10% glycerol) were resolved on 4–20% Criterion TGX Stain-Free Precast Polyacrylamide Gels (Bio-Rad), transferred to nitrocellulose membranes (Bio-Rad), and incubated for 1 hr in blocking buffer (5% non-fat dry milk in 0.1% TBS-Tween) followed by overnight incubation with anti-Trio monoclonal antibody (1:1000; DHSB #9.4A) diluted in blocking buffer. Primary antibody was detected using species-specific horse-radish peroxidase (HRP) conjugated secondary antibody (Jackson ImmunoResearch) with enhanced chemiluminescence (ECL,Sigma). Densitometry analysis was performed using Image Lab software (Bio-Rad), and significance was calculated using group analysis on GraphPad (Prism)

#### *Drosophila* brain dissection, immunohistochemistry, visualization, and statistical analysis

For *Drosophila* mushroom body morphology imaging, brains were dissected using #5 Dumont fine forceps (Ted Pella, Inc.) in PBS supplemented with 0.1% Triton X-100 (0.1% PBS-T). The proboscis was removed to provide a forceps grip point, and the remaining cuticle and trachea were peeled away from the brain. Brains were submerged in 1X PBS on ice until all brains were dissected. Dissected brains were fixed in 4% paraformaldehyde for 30 min and then permeabilized in 0.3% PBS-Triton X-100 (0.3% PBS-T) for 30 min, on ice. Brains were carefully transferred to 0.5 ml Eppendorf tubes in 0.1% PBS-T. For both primary and secondary antibody incubations, brains were left rocking at 4°C for 24-72 hours (see list of dilutions and incubation times below) in 0.1% PBS-T supplemented with normal goat serum (Jackson ImmunoResearch) at a 1:20 dilution. Immunostained brains were mounted on SuperFrost Plus slides in Vectasheild (Vector Laboratories) using a coverslip bridge. Brains were imaged on a Nikon A1R confocal microscope. Maximum intensity projections were generated using Fiji ImageJ software.

**Antibody:** Mouse monoclonal 9.4A anti-Trio, **Incubation time:** 48-72hr, **Dilution:** 1:50

**Antibody:** Mouse monoclonal 1D4 anti-Fasciclin II, **Incubation time:** 48-72hr, **Dilution:** 1:50

**Antibody:** Rabbit Polyclonal Anti-Green Fluorescent Protein (GFP), **Incubation time:** Overnight-24hr, **Dilution:** 0.125:100

**Antibody:** Alexa Fluor® 488 AffiniPure Polyclonal Goat Anti-rabbit IgG, **Incubation time:** Overnight-24hr, **Dilution:** 1:100

**Antibody:** Cy™3 AffiniPure Polyclonal Goat Anti-Mouse IgG, **Incubation time:** Overnight-24hr, **Dilution:** 1:100

#### *Drosophila* neuron live imaging confocal microscopy, neuronal reconstruction, data analysis, and statistical analysis

Live imaging of class IV ddaC neurons was performed as described previously (Iyer *et al*., 2012). Briefly, wondering 3^rd^ instar *ppk-Gal4,UAS-mcd8::GFP* labeled larvae were mounted in 1:5 (v/v) diethyl ether: halocarbon oil under an imaging bridge of 22 x 22mm glass coverslips topped with a 22 x 55mm glass coverslip. The ddaC images were captured on a Nikon A1R inverted Confocal microscope. Maximum intensity projections were generated using Fiji ImageJ software. Quantitative morphological data were compiled using the Simple Neurite tracer (SNT) plugin for Fiji (Ferreira *et al*., 2014; Arshadi *et al*., 2021). Batch processing was completed using a custom Fiji macro and Rstudio script created and gifted by Dr. Atit A. Patel (Dr. Dan Cox Lab, Georgia State University) (Lottes *et al*., 2023) and the resulting data were exported to Excel (Microsoft).

### Statistical Analysis

Group analysis on biological triplicate experiments was performed using One-Way or Two-Way ANOVA (Tukey’s multiple-comparison test) on GraphPad (Prism). Sample sizes (n) and p-values are denoted in text, figures, and/or figure legends and indicated by asterisks (e.g., *p<0.05).

## Acknowledgements

This work was supported by the Emory University Emory Integrated Cellular Imaging Core Facility (RRID:SCR_023534). The authors thank Dr. Dan Cox, GA State Neuroscience Institute, and his lab members Drs. Erin Lottes and Atit Patel for training, reagents, and discussions. We also thank the members of the Moberg and Corbett laboratories, especially Dr. Milo Fasken for insightful discussions.

## Abbreviations

RBP: RNA binding protein
Pab: polyadenosine binding protein
ZC3H14: zinc finger Cys-Cys-Cys His type containing 14
Nab2: nuclear polyadenosine RNA binding protein 2
GEF: guanine nucleotide exchange factor

## Supplemental Figure Legends

**Supplemental Figure 1.** Trio is present in *Drosophila* Kenyon cell bodies and Caylx. (**A**) Diagram of the adult *Drosophila* mushroom body depicting the Caylx, peduncle, protocerebellar bridge, fan-shaped body, ellipsoid body, and axons of the vertically projecting alpha (α) and alpha prime (α’) neurons, the medially projecting gamma (γ), beta (β), and beta prime (β’) neurons. (**B**) Immunofluorescence images of Control (*w-;UAS-mcd8::GFP/201Y-Gal4;;*) mushroom bodies driving *UAS-mcd8::GFP* under the α, β, γ lobe-specific mushroom body *201Y-Gal4* driver. (**C**) Immunofluorescence images of Control (*w-;UAS-mcd8::GFP/C305a-Gal4;;*) mushroom bodies driving *UAS-mcd8::GFP* under the α’ and β’ lobe-specific mushroom body *C305a-Gal4* driver. False colored panels show fluorescence corresponding to α-GFP (green, mcd8::GFP), α-trio (purple), and merges of the channels. Scale bar = 50 μm.

**Supplemental Figure 2.** Heterozygosity for Mettl3 does not dominantly rescue *Nab2^null^* mushroom body phenotypes. (**A**) Representative max projections of mushroom bodies of indicated genotypes stained by α-FasII. (**B**) Quantification of frequency of α lobe defects in each indicated genotype. (**C**) Quantification of frequency of β lobe defects in indicated genotypes. Scale bar = 50 μm.

**Supplemental Figure 3.** Expression of Rac1 or Rho1/A in the mushroom body of *Nab2^null^* flies is lethal. (**A**) Percent of Control, *Nab2^null^*, *UAS-Rho1, UAS-Rac1, UAS-Rho1;Nab2^null^* and *UAS-Rac1;Nab2^null^* flies that eclose as viable adults (calculated as #observed/#expected) using the *OK107-Gal4* mushroom body driver.

