## Supplementary figures and images for "The RNA binding protein Nab2 regulates splicing of the RhoGEF *trio* transcript to govern axon and dendrite morphology"

### Supplemental Figure 1

Supplemental Figure 1
Lancaster et al.

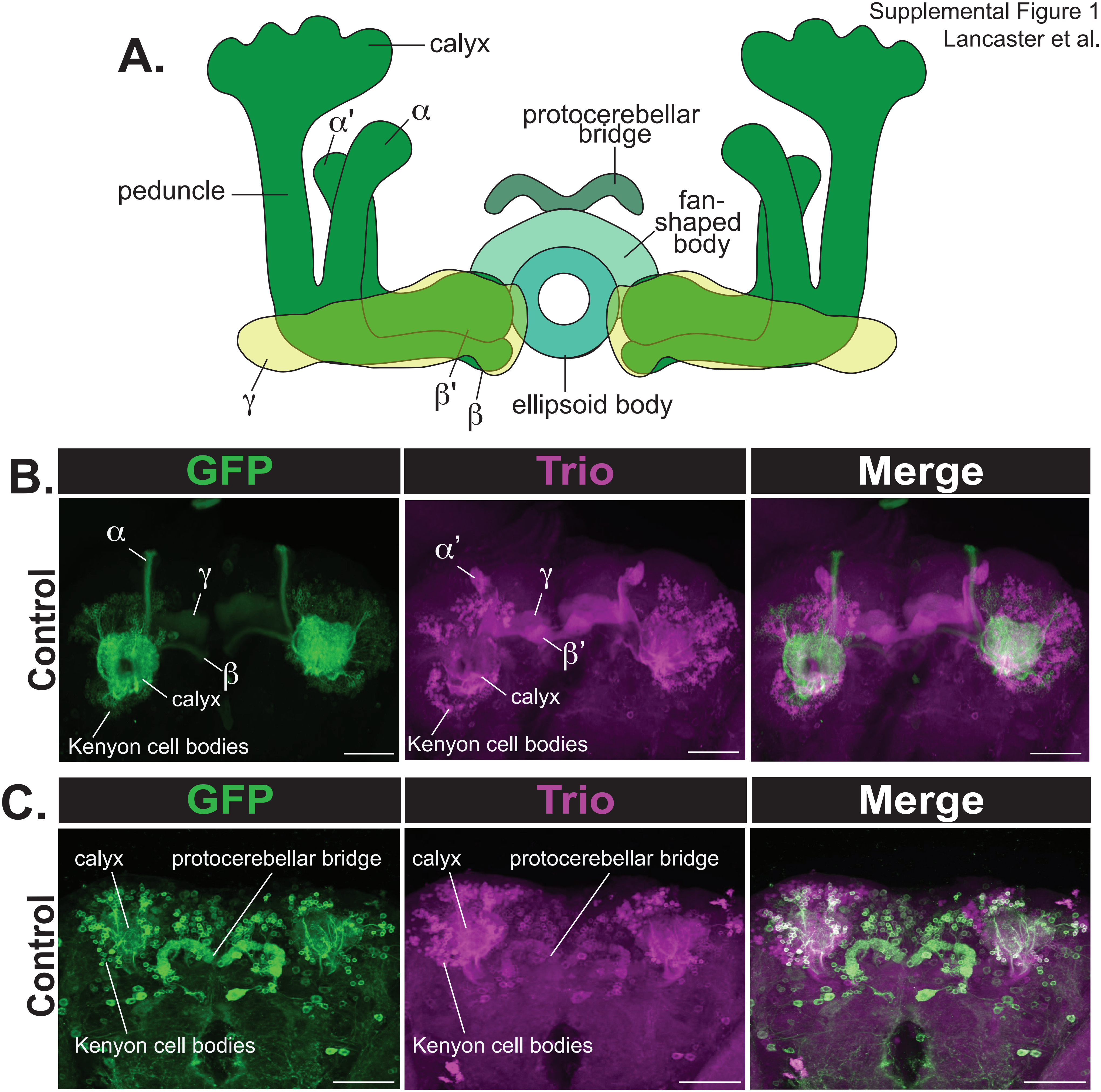

### Supplemental Figure 2

Supplemental Figure 2
Lancaster et al.

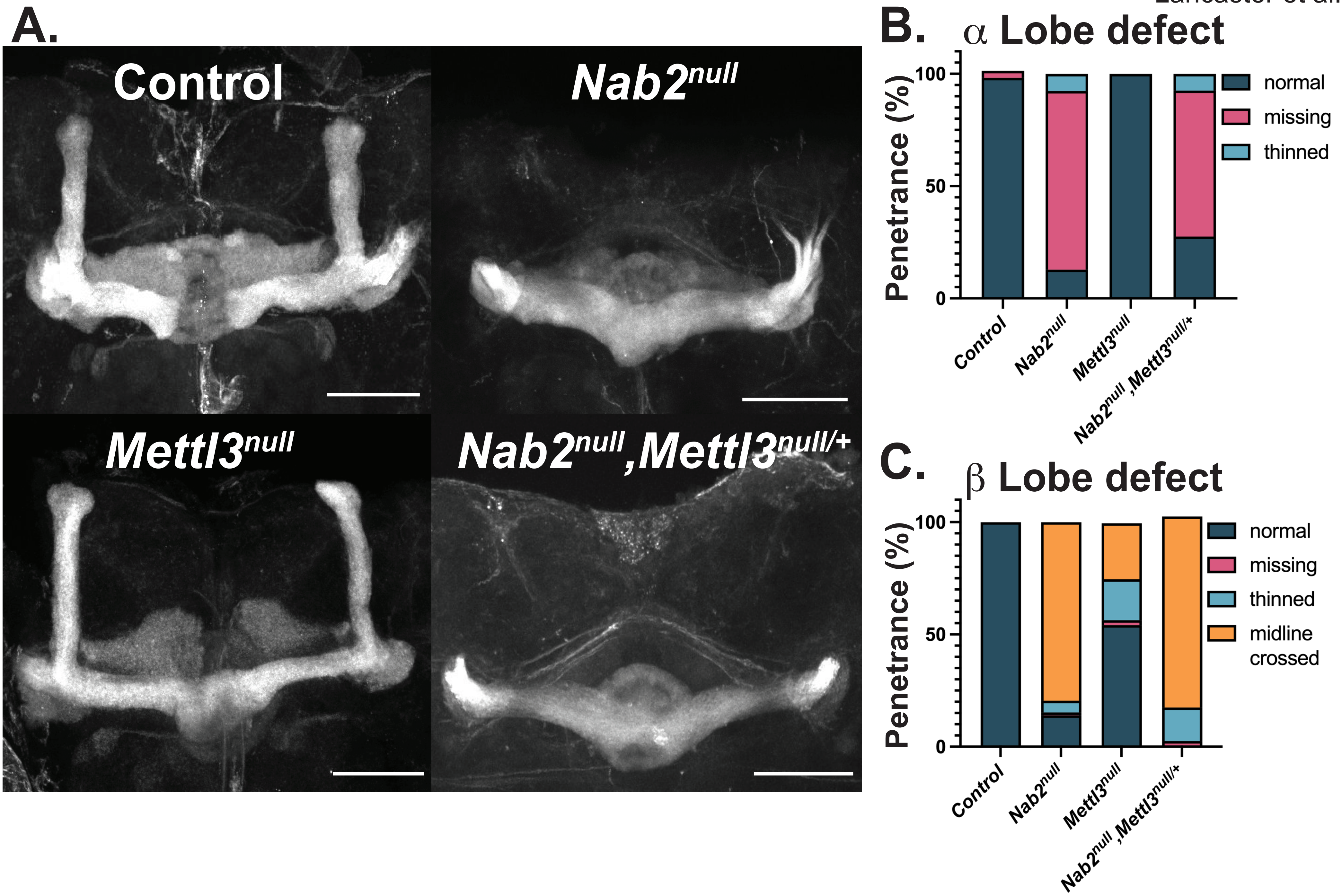

### Supplemental Figure 3

Supplemental Figure 3
Lancaster et al.

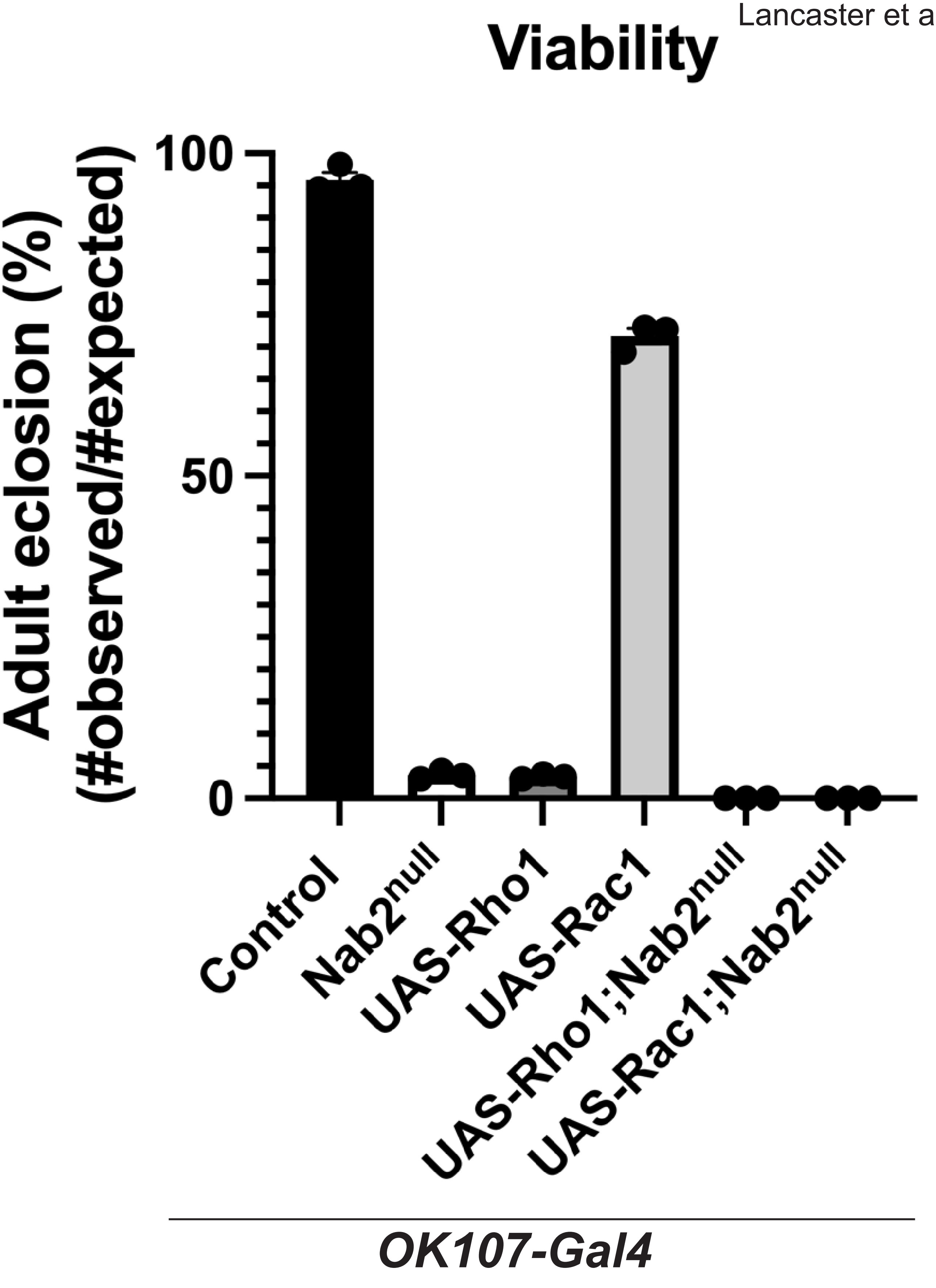
